# Complementary α-arrestin - Rsp5 ubiquitin ligase complexes control selective nutrient transporter endocytosis in response to amino acid availability

**DOI:** 10.1101/2020.04.24.059832

**Authors:** Vasyl Ivashov, Johannes Zimmer, Sinead Schwabl, Jennifer Kahlhofer, Sabine Weys, Ronald Gstir, Thomas Jakschitz, Leopold Kremser, Günther K. Bonn, Herbert Lindner, Lukas A. Huber, Sébastien Léon, Oliver Schmidt, David Teis

## Abstract

How cells adjust transport across their membranes is incompletely understood. Previously, we have shown that *S.cerevisiae* broadly re-configures the nutrient transporters at the plasma membrane in response to amino acid availability, through selective endocytosis of sugar- and amino acid transporters (AATs) (Müller et al., 2015). A genome-wide screen now revealed that Art2/Ecm21, a member of the α-arrestin family of Rsp5 ubiquitin ligase adaptors, is required for the simultaneous endocytosis of four AATs and induced during starvation by the general amino acid control pathway. Art2 uses a basic patch to recognize C-terminal acidic sorting motifs in these AATs and instructs Rsp5 to ubiquitinate proximal lysine residues. In response to amino acid excess, Rsp5 instead uses TORC1-activated Art1 to detect N-terminal acidic sorting motifs within the same AATs, which initiates exclusive substrate-induced endocytosis of individual AATs. Thus, amino acid availability activates complementary α-arrestin-Rsp5-complexes to control selective endocytosis for nutrient acquisition.

## Introduction

Cells regulate the import of amino acids, glucose, and other nutrients to fuel metabolism, sustain growth or maintain homeostasis. For the selective transport of amino acids across the plasma membrane (PM) and other cellular membranes, the human genome encodes more than 60 known amino acid transporters (AATs). Mutations in AATs cause severe defects of amino acid metabolism, and the deregulation of AATs is linked to a range of human pathologies, including neurodegenerative diseases, diabetes and cancer (Kandasamy et al., 2018, Smith, 1990, McCracken and Edinger, 2013, Zhang et al., 2017).

AATs belong to the family of solute carriers (SLCs) and form selective pores that change from an outward- to an inward-facing conformation and thereby transport their amino acid substrates across membranes. Hence, the addition of AATs to the PM or their selective removal by endocytosis determines quantity and quality of amino acid transport. In human cells the molecular mechanism leading to the selective endocytosis of AATs are largely unclear.

The mechanisms for the endocytic down-regulation of AATs are beginning to emerge in the budding yeast*, S. cerevisiae*. Yeast cells frequently experience acute fluctuations in amino acid availability (Broach, 2012) and can rapidly remodel their 24-26 AATs to optimize the import of amino acids with regard to the quantity and quality of the nitrogen source available (Grenson et al., 1966, Grenson, 1966, Andre, 2018, Van Belle and Andre, 2001). The selective, ubiquitin-dependent endocytosis of AATs and other integral PM proteins frequently requires the HECT-type ubiquitin ligase Rsp5, the orthologue of Nedd4 in humans (Hein et al., 1995, Dupré et al., 2004, Belgareh-Touze et al., 2008). To specifically recognize and ubiquitinate PM proteins, Rsp5 interacts with a family of ubiquitin ligase adaptors, the α-arrestin proteins (Nikko et al., 2008, Lin et al., 2008). The yeast genome encodes 14 α-arrestin proteins (arrestin-related trafficking adaptors, ARTs; Art1-10, Bul1-3 and Spo23), which function in a partially redundant manner (reviewed in (Becuwe et al., 2012a, MacGurn et al., 2012, Babst, 2020, O’Donnell and Schmidt, 2019). Six α-arrestins have been identified in humans (Alvarez, 2008). α-Arrestin proteins use PPxY (PY) motifs to interact directly with the three WW domains of Rsp5/Nedd4 (Becuwe et al., 2012a, Rauch and Martin-Serrano, 2011). How α-arrestins recognize their cargoes in general, and more specifically AATs, is only partially understood, but appears to involve their arrestin domains. These domains have large, disordered loop- and tail-like insertions that contribute to the function of ARTs (Baile et al., 2019).

The activity of α*−*arrestins is controlled by complex post-translational modifications (PTMs) involving (de-)ubiquitination and (de-)phosphorylation. In several cases the binding of α*−*arrestins to Rsp5 results in their own ubiquitination, which is required for function (Lin et al., 2008, Becuwe et al., 2012b, Hovsepian et al., 2017). In case of Art1 ubiquitination determines its localization (Lin et al., 2008). Additional ubiquitination of ARTs can target them for proteasomal degradation, which is counteracted by deubiquitinating enzymes (Ho et al., 2017). Several α*−*arrestins are activated by dephosphorylation events in response to nutrient availability or cellular stress (Becuwe et al., 2012a, O’Donnell et al., 2013, Alvaro et al., 2014, Becuwe et al., 2012b, Becuwe and Leon, 2014, Merhi and André, 2012). One example are high levels of amino acids, which activate the target of rapamycin complex 1 (TORC1). This leads to inhibition of the kinase Npr1 and thereby promotes the activity of Art1 (MacGurn et al., 2011). At the PM, Ppz phosphatases activate Art1 by dephosphorylation of its Npr1-dependent phosphorylation sites (Lee et al., 2019) and thereby stimulate ubiquitination and endocytosis of the Art1 cargoes Mup1 (methionine transporter), Can1 (arginine transporter), Tat2 (tyrosine and tryptophan transporter) and Lyp1 (lysine transporter) in response to substrate excess. Similarly, TORC1 signaling also promotes endocytosis of the general amino acid permease (Gap1) via the ARTs Bul1/2 (Merhi and Andre, 2012). It seems that nutrient-dependent TORC1 activation controls α*−*arrestin-mediated ubiquitin-dependent AAT endocytosis to adjust amino acid influx to metabolic needs.

Activated Art1 recognizes specific sorting signals in AATs. The flux of methionine through Mup1 or arginine through Can1 requires a conformational change into the inward-facing conformation. This conformational change probably dislodges a C-terminal plug, that otherwise ‘seals’ the pore and at the same time exposes an N-terminal acidic patch that is recognized by activated Art1. Art1 interacts with this acidic patch and then orients Rsp5 to ubiquitinate nearby lysine residues (Guiney et al., 2016, Gournas et al., 2018, Busto et al., 2018, Ghaddar et al., 2014). Similar results have been obtained for the uracil transporter Fur4 (Keener and Babst, 2013). A further layer of complexity may be added by stimulus-induced phosphorylation of nutrient transporters (Marchal et al., 1998, Paiva et al., 2009), akin to the recognition of G-protein-coupled receptors (GPCRs) by β–arrestins in mammalian cells (Nikko et al., 2008, Nobles et al., 2011). Hence, a molecular basis of α*−*arrestin-dependent nutrient transporter endocytosis in response to excess nutrients is emerging. We refer to this process as substrate-induced endocytosis.

Yet, not only substrate transport can induce nutrient transporter endocytosis, but also nutrient limitation, acute starvation or rapamycin treatment (Crapeau et al., 2014; Hovsepian et al., 2017; Jones et al., 2012; Laidlaw et al., 2020; Lang et al., 2014; Müller et al., 2015; Schmidt et al., 1998; Yang et al., 2020). Starvation-induced endocytosis is part of a coordinated catabolic cascade. Together with proteasomal degradation and with macro- and micro-autophagy it maintains amino acid homeostasis to allow entry into quiescence for cell survival during nutrient limitation. This catabolic cascade at the onset of starvation appears to be conserved in metazoans (Mejlvang et al., 2018, Edinger and Thompson, 2002a, Edinger and Thompson, 2002b, Vabulas and Hartl, 2005, Suraweera et al., 2012).

The molecular mechanisms of starvation-induced endocytosis are not clear. Hence, there is a significant knowledge gap in understanding how cells control nutrient transporter abundance upon amino acid and nitrogen scarcity. Here, we have characterized systematically how amino acid abundance alters the nutrient transporter repertoire at the PM of the budding yeast *S. cerevisiae*. We find that TORC1 signaling and the general amino acid control (GAAC) pathway toggle Art1- or Art2-Rsp5 complex activities to induce endocytosis of the same set of four different AAT depending on amino acid availability. In these AATs, activated Art1- or Art2-Rsp5 complexes recognize distinct acidic sorting signals in the N-terminal (Art1 sorting signal) or C-terminal (Art2 sorting signal) regions to initiate Rsp5-dependent ubiquitination and subsequent endocytic down-regulation. Using such complementary α*−*arrestins switches in combination with distinct acidic sorting motifs in a single nutrient transporter could be a more general mechanism to adjust transport across cellular membranes and thereby to meet metabolic demands.

## Results

### Amino acid availability induces selective endocytosis of nutrient transporters

We employed *S. cerevisiae* as a model system to address how eukaryotic cells adjust their nutrient transporters at the plasma membrane (PM) to nutrient availability. First, we used live cell fluorescence microscopy to analyze in yeast cells the localization of 147 putative PM proteins that were C-terminally GFP-tagged at their native chromosomal locus (Saier et al., 2016, Babu et al., 2012, Breker et al., 2014). In cells growing exponentially under defined (rich) conditions we detected 49 GFP-tagged proteins at the PM, including seven different amino acid transporters (AATs) and six different carbohydrate transporters (Fig. 1A, S1A, Table S1). A fraction of these proteins was additionally detected inside the vacuole (Fig. 1B, S1A), suggesting continuous turnover.

**Figure 1:**
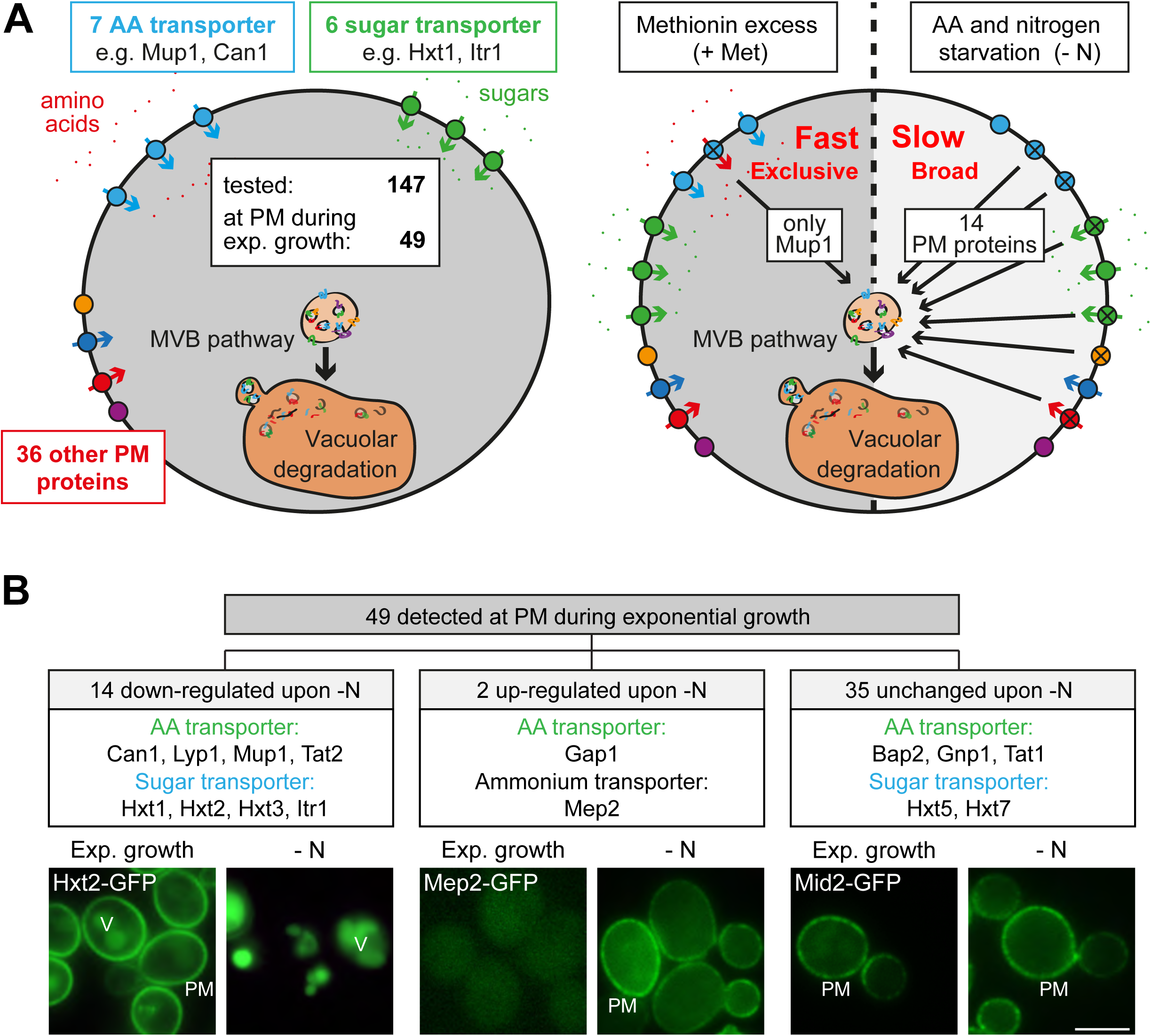
Amino acid and nitrogen starvation triggers broad but specific endocytosis and lysosomal degradation of plasma membrane proteins. A) Left: a library of 147 yeast strains expressing chromosomally GFP-tagged membrane proteins was tested for plasma membrane (PM) localization during nutrient replete exponential growth. Right: verified PM proteins were starved for amino acids and nitrogen (- N) 6-8h or treated with 20 µg/ml L-methionine (+Met) after 24h of exponential growth. The localization of GFP was assayed by fluorescence microscopy. B) Summary of the phenotypes of GFP-tagged PM proteins during starvation. Indicated are numbers of PM proteins that are down-regulated, up-regulated or unchanged compared to the exponential growth phase, each exemplified by one representative strain. PM: plasma membrane; V: vacuole. Scale bars = 5 µm. See also Fig. S1 and Table S1.

Next, we examined how amino acid availability affected this repertoire of PM proteins and used initially the high affinity methionine transporter Mup1 as a model cargo because its regulation in response to nutrient excess is well characterized (Busto et al., 2018, Lee et al., 2019, Gournas et al., 2018, Guiney et al., 2016, Baile et al., 2019). Moreover, Mup1 is one of the most abundant PM proteins and it is easy to follow its endocytosis and subsequent transport into the vacuole (Busto et al., 2018). In absence of methionine in the growth medium Mup1-GFP localized to the PM (Fig. S1B). Low levels of methionine in the growth medium did not efficiently trigger its endocytosis (Fig. S1B). Yet, once a critical methionine concentration in the growth medium was reached, the vast majority of Mup1-GFP was removed from the PM by endocytosis and was subsequently transported to the vacuole via the multivesicular body (MVB) pathway (Fig. S1B). Substrate-induced endocytosis was rapid and efficient: within 60-90 minutes after methionine addition Mup1 was almost quantitatively delivered to the vacuole. Excess methionine appeared to exclusively induce endocytosis of Mup1 since all other tested PM proteins remained at the cell surface (Fig. 1A, Table S1). This exclusive substrate-induced endocytosis of Mup1 was also dependent on its methionine transport activity as shown by using the Mup1-G78N mutant, which cannot transition to the open-inward conformation (Busto et al., 2018) (Fig. S1C). These results are consistent with earlier reports showing that the high-affinity AATs Mup1, Can1, Tat2 and Lyp1 undergo exclusive endocytic down-regulation in response to excess of their respective substrates but remained stable at the PM when the substrate of another permease was present in excess (Gournas et al., 2017, Busto et al., 2018, Nikko and Pelham, 2009).

Others and we had shown earlier that amino acid and nitrogen starvation (hereafter starvation) induced the degradation of membrane proteins via the MVB pathway (Müller et al., 2015, Jones et al., 2012). Consistently, in response to starvation 14 (out of 49) different PM proteins were selectively removed from the PM and transported into the vacuole, including four AATs (Mup1, Can1, Lyp1, Tat2) and four carbohydrate transporters (Hxt1, Hxt2, Hxt3, Itr1) (Fig. 1B, Table S1). PM localization of the general amino acid permease Gap1 and the ammonium permease Mep2 was up-regulated, and the localization of 35 GFP-tagged proteins to the PM remained largely unchanged, although in some instances the vacuolar GFP fluorescence was increased (Fig. 1A, B, S1A, Table S1). The starvation-induced endocytosis of Mup1 was independent of its capability to transport its substrate (Fig. S1C), and was also observed upon starvation for individual essential amino acids (Fig. S1D). We re-evaluated the starvation-induced endocytosis of Mup1, Can1, Lyp1, Tat2, Fur4, Hxt1, Hxt2 and Hxt3 in a different genetic background (SEY6210; Fig. S1B, 3C, S3B, S6D). The overall response was similar.

Hence, in response to starvation, a broad range of nutrient transporters was selectively removed from the PM by endocytosis and transported into the vacuole within 3-6 hours. In contrast, substrate-induced endocytosis of AATs was quick and exclusive (Fig. 1A). Earlier work showed that substrate-induced endocytosis of the AATs Mup1, Can1 and Lyp1 required TORC1 signaling to allow ubiquitination of the transporters by the HECT-type ubiquitin ligase Rsp5 in complex with the α-arrestin Art1 (Lin et al., 2008, Guiney et al., 2016, Busto et al., 2018, Gournas et al., 2018, MacGurn et al., 2011). Remarkably, the same set of AATs underwent simultaneous starvation-induced endocytosis, but the molecular mechanism was not clear.

### A genome-wide screen identifies regulators of starvation-induced endocytosis of AATs

To identify the genes that are required for the starvation-induced endocytosis of Mup1, we conducted a fluorescence-based genome-wide screen. We measured Mup1 sorting into the lumen of vacuoles via the MVB pathway by fusing the pH-sensitive GFP variant pHluorin to the C-terminus of Mup1 (Mup1-pHluorin) (Henne et al., 2015, Prosser et al., 2010). Fluorescence microscopy and fluorescence-activated cell sorting (FACS) (Fig.2A, B) showed a strong signal of Mup1-pHluorin at the PM of cells growing under defined rich conditions. In response to starvation the fluorescence of Mup1-pHluorin was efficiently quenched in the acidic vacuoles of wild type (WT) cells, but not when the MVB pathway was blocked (e.g. ESCRT-I mutant *vps23Δ*) (Fig. 2A, B). Hence Mup1-pHluorin is a suitable reporter to identify mutants that block trafficking of Mup1 from the PM into the vacuole.

**Figure 2:**
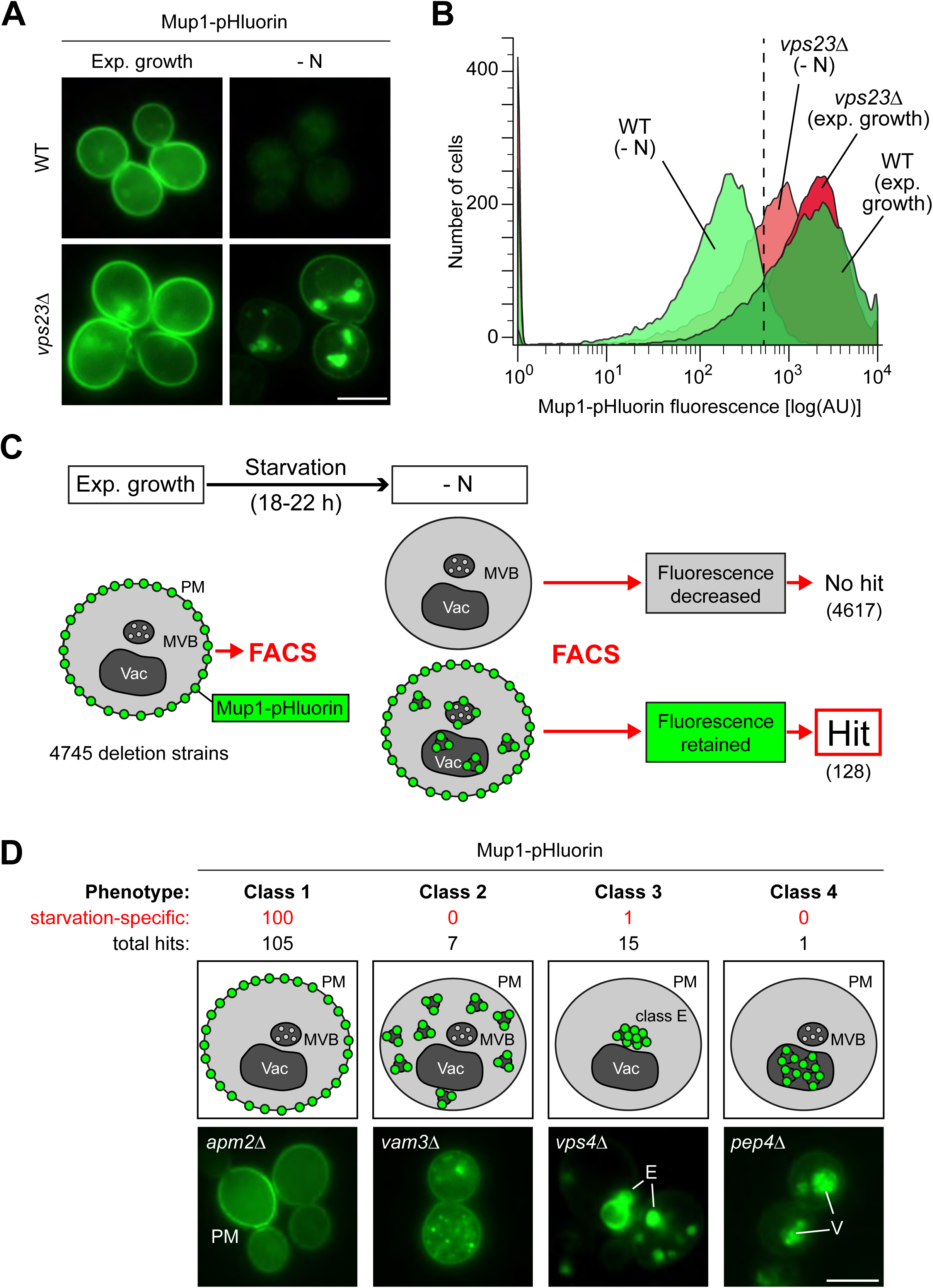
A genome wide screen revealed genes affecting Mup1-pHluorin endocytosis during starvation. **A)** Live-cell fluorescence microscopy analysis of WT (BY4742) and *vps23Δ* cells expressing *MUP1-pHluorin* from plasmid and starved (- N) for 18-22h. The images exemplify quenched pHluorin fluorescence in vacuoles of wild type (WT)-like cells and retained fluorescence in mutants with defects in the starvation-induced endocytosis of Mup1-pHluorin. **B)** The strains from (A) were exponentially grown in 96-well plates for 5h and starved (- N) for 18-22h. At least 15000 cells from each strain and condition were analyzed by flow cytometry. The exemplified histograms display decrease of fluorescence in wild type (WT)-like strains and fluorescence retention in mutants with defects in the starvation-induced endocytosis of Mup1-pHluorin (e.g. *vps23*Δ). **C)** Workflow of the flow-cytometry-based genome-wide screen for mutants defective in starvation-induced endocytosis of Mup1-pHluorin. **D)** Summary of phenotypes of all mutants scored in the starvation-induced endocytosis screen (C) as determined by fluorescence microscopy. Class 1 mutants retain Mup1-pHluorin fluorescence at the plasma membrane (PM); class 2 mutants in small cytosolic objects; class 3 mutants in class E-like objects (E); class 4 mutants within vacuoles (V). Each phenotype is exemplified by one representative deletion mutant. Indicated are the numbers of strains that share a similar phenotype and the number of hits specific for starvation-induced endocytosis of Mup1-pHluorin (red). Scale bars = 5 µm. See also Fig. S2 and table S2.

Next, Mup1-pHluorin was introduced into the yeast non-essential knock-out collection. 4745 mutants expressing Mup1-pHluorin were grown in selection medium to exponential phase before they were subjected to starvation for 18-22 hours (Fig. 2C). Automated FACS was used to measure Mup1-pHluorin fluorescence intensity of at least 15000 cells before and after starvation. The vast majority of mutants efficiently quenched the fluorescence of Mup1-pHluorin after starvation, but 128 mutants still exhibited Mup1-pHluorin fluorescence after starvation (Fig. 2C, Table S2). To determine at which step Mup1-pHluorin transport into vacuoles was blocked in these mutants, we used live cell fluorescence microscopy. This allowed us to classify four phenotypes (Fig 2D, Table S2). Class 1 mutants retained Mup1-pHluorin at the PM after starvation (105 mutants, e.g. *apm2Δ* mutants). In class 2 mutants Mup1-pHluorin was detected on small intracellular objects (7 mutants, e.g. *vam3Δ* mutants). Class 3 mutants accumulated Mup1-pHluorin in larger class E compartment-like objects (15 mutants, e.g. *vps4Δ* mutants). One Class 4 mutant (*pep4Δ*) did not quench Mup1-pHluorin fluorescence inside the vacuole, as it is deficient in the main lysosomal protease. Gene ontology (GO) analysis confirmed that our screen identified major general regulators of endosomal transport (e.g. the ESCRT complexes) and was enriched for proteins that are annotated as components of the PM, endosomes and vacuoles (Fig. S2, Table S3) suggesting that is was successful. In addition, we identified many genes and components of protein complexes that would not be predicted to be involved in the regulation of endocytosis (Fig. S2, Table S2, S3).

Next, we aimed to narrow down genes that function specifically during starvation-induced endocytosis and to distinguish them from general endocytic regulators (i.e. core components of endocytic trafficking machineries). Therefore, most mutants (124) were exposed to methionine excess for 90 minutes and then subjected to live cell fluorescence microscopy. Of the mutants falling into classes 2 - 4, the majority was required for Mup1 sorting into the vacuole in response to both starvation and methionine excess. Among them were many mutants in subunits of well-characterized protein complexes that constituted the core machinery of endo-lysosomal trafficking (Fig. S2, Table S2, S3). However, the majority (100) of the class 1 mutants (Mup1-pHluorin retained at the PM) was specifically required for starvation-induced endocytosis but not for methionine-induced endocytosis (Fig. 2D, Table S2).

### The α−arrestin Art2 is specifically required for amino acid transporter endocytosis in response to starvation

A key step for the endocytic down-regulation of PM proteins is their selective ubiquitination. This selectivity is mediated by α*−*arrestin molecules that direct the HECT-type ubiquitin ligase Rsp5 to ubiquitinate specific PM proteins (Babst, 2020). Among the 14 α*−*arrestins encoded by the yeast genome, our screen identified only Ecm21/Art2 to be specifically required for starvation-induced endocytosis of Mup1.

We compared Art2-dependent starvation-induced endocytosis to Art1-dependent substrate-induced endocytosis of Mup1-GFP. Efficient methionine-induced endocytosis of Mup1 required Art1 but not Art2 (Fig. 3A), consistent with earlier observations (Guiney et al., 2016, MacGurn et al., 2011, Busto et al., 2018). Starvation-induced endocytosis of Mup1 was slower than methionine-induced endocytosis. In WT cells, 240 – 360 minutes after onset of starvation the majority of Mup1-GFP was removed from the PM and sorted into vacuoles for degradation (Fig. 3A, B). In *art2Δ* mutants, but not in *art1Δ* mutants, the vast majority of Mup1-GFP remained at the PM and was no longer delivered to vacuoles (Fig. 3A). This was confirmed by Western blot (WB) analysis from total cell lysates. In WT cells, the proteolytic degradation of Mup1-GFP inside vacuoles released free GFP, which remained stable and thus could be monitored by Western blotting (Fig. 3B, lane 2-4). In *art2Δ* mutants only little free GFP could be detected (Fig. 3B lane 7-8). Re-expression of Art1 or Art2 in the corresponding mutants restored methionine- or starvation-induced Mup1 endocytosis, respectively (Fig. S3A). Importantly, Art2 was required for the efficient starvation-induced endocytosis of all four AATs, Mup1, Can1, Lyp1, and Tat2 (Fig. 3A, C). Interestingly, the same four AATs (Mup1, Can1, Lyp1 and Tat2) also depend on Art1 for efficient substrate-induced endocytosis (Gournas et al., 2017, MacGurn et al., 2011, Lin et al., 2008). Starvation-induced endocytosis of other PM proteins (e.g. the uracil transporter Fur4 or the hexose transporters Hxt1 and Hxt2) was largely independent of Art2 (Fig. S3B).

**Figure 3:**
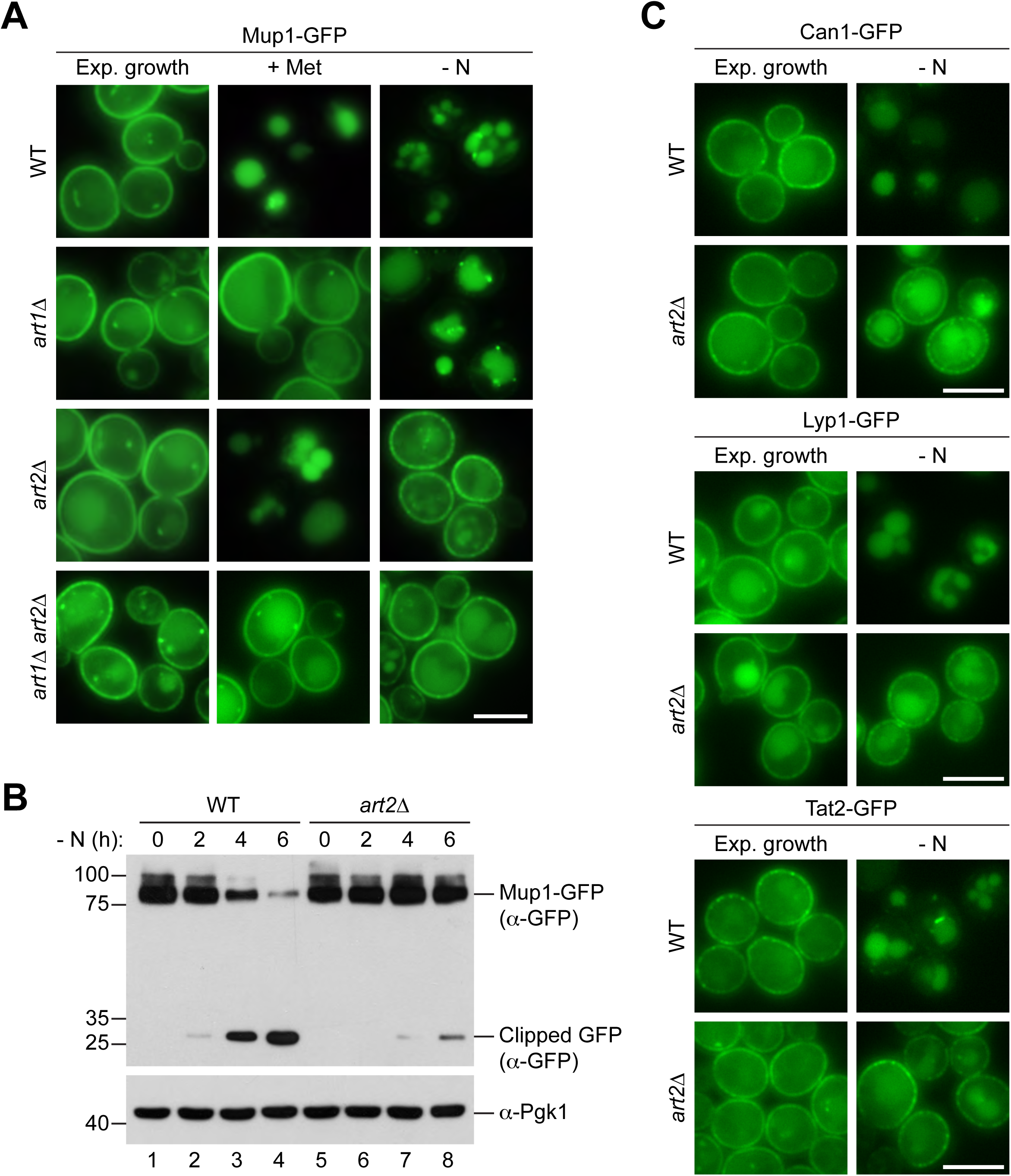
Art1 and Art2 are non-redundant in promoting substrate- and starvation-induced endocytosis of amino acid transporters. **A)** Live-cell fluorescence microscopy analysis of Mup1-GFP endocytosis in wild type (WT), *art1*Δ, *art2*Δ and *art1*Δ *art2*Δ cells expressing *MUP1-GFP* from plasmid. Cells were treated with 20 µg/ml L- methionine (+ Met) for 1.5h or starved (- N) for 6h after 24h exponential growth. **B)** SDS PAGE and Western blot analysis with the indicated antibodies of whole cell protein extracts from wild type (WT) and *art2*Δ cells expressing *MUP1-GFP* that were starved (- N) for the indicated times after 24h exponential growth. **C)** Live-cell fluorescence microscopy analysis of wild type (WT) and *art2*Δ cells expressing *CAN1-GFP*, *LYP1-GFP* or *TAT2-GFP*. Cells were starved (- N) for 6h after 24h exponential growth. Scale bars = 5 µm. See also Fig. S3.

These results suggested a swap between the α*−*arrestins Art1 and Art2 by amino acid availability as a rule for the regulation of four AATs. The starvation-induced endocytosis of Mup1, Can, Lyp1 and Tat2 required Art2. Substrate-induced endocytosis of Mup1, Can1 and Lyp1 required Art1, whereas in case of Tat2 substrate-induced endocytosis was less stringent and could be mediated either by Art1 or Art2 (Nikko and Pelham, 2009).

### Up-regulation of Art2 by the GAAC pathway drives starvation-induced endocytosis of Mup1

Our experiments so far demonstrated that Art1 and Art2 mediate the endocytic down-regulation of a set of AATs in response to changes in amino acid availability in a mutually exclusive manner. Hence their activity must be strictly regulated. α*−*arrestin proteins are subject to complex multi-level regulation and their activation can require transcriptional regulation, (de-)phosphorylation and ubiquitination (Hovsepian et al., 2017, Lin et al., 2008, Becuwe et al., 2012b). WB analysis of total cell lysates indicated quantitative band shifts of Art1 in response to starvation (Fig. 4A, lanes 2,3) that most probably represented Npr1-dependent phosphorylation and thus inhibition of Art1 activity upon TORC1 inactivation as previously described (MacGurn et al., 2011). The protein levels of the functional 6xHis-TEV-3xFlag-tagged 170kD protein Art2-HTF (Fig. S4A) were comparably low under rich conditions (Fig. 4A, lane 4, Fig. S4A), but were up-regulated in response to starvation (Fig. 4A, lanes 5,6). Quantitative reverse transcription (RT) PCR analysis indicated that also the *ART2* mRNA levels increased during starvation (Fig. 4B).

**Figure 4:**
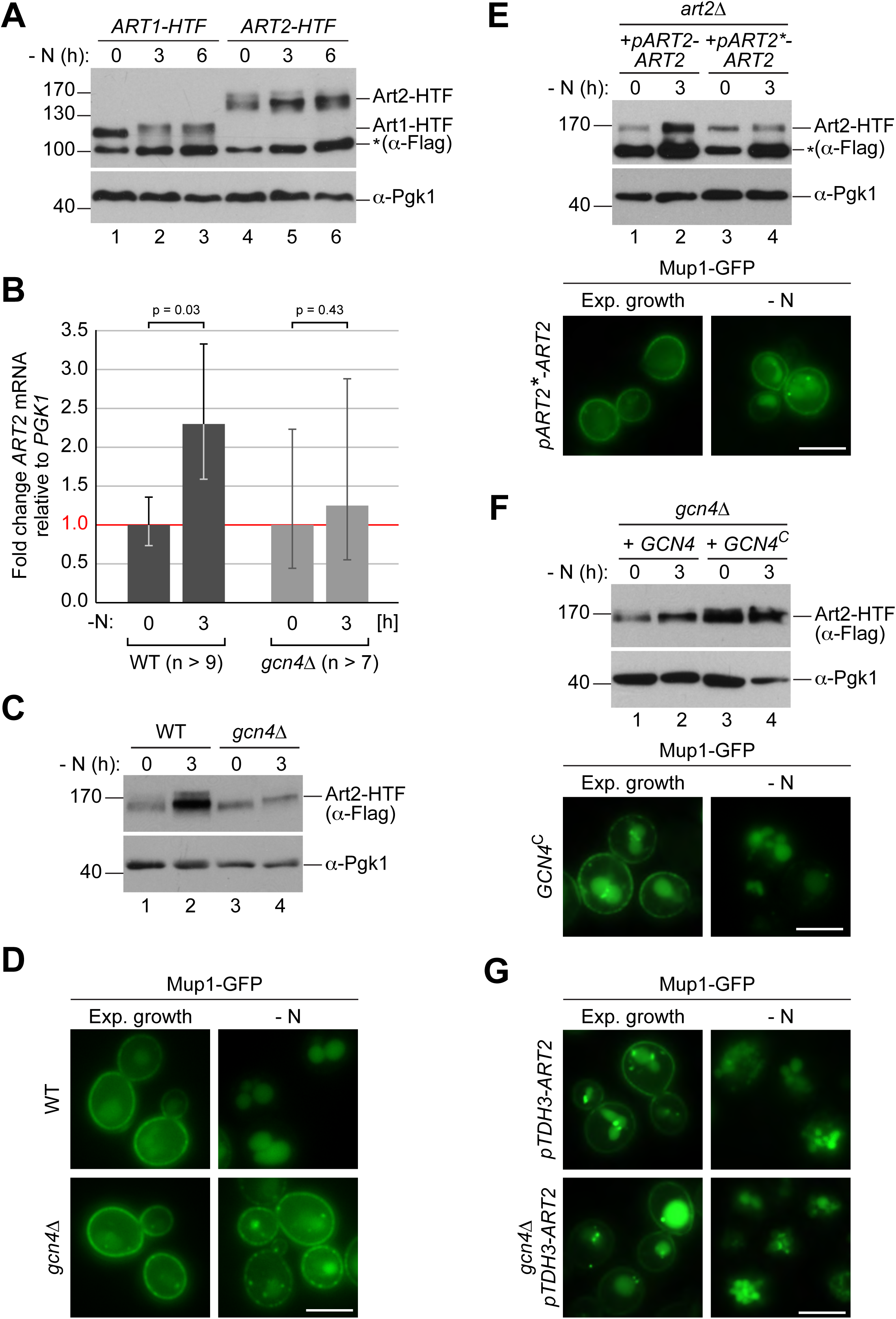
The general amino acid control pathway promotes starvation-induced endocytosis Mup1 by up-regulating Art2. **A)** SDS PAGE and Western blot analysis with the indicated antibodies of whole cell protein extracts from WT cells expressing *ART1-HTF* or *ART2-HTF*. Cells were starved (- N) for the indicated times after 24h exponential growth. The asterisk indicates a non-specific background band of the FLAG antibody. **B)** RT-qPCR analysis of *ART2* transcript levels (normalized to the stable *PGK1* transcript) in wild type (WT) and *gcn4*Δ cells. Cells were starved (- N) for 3h after 24h exponential growth. Values are presented as fold-change of the starting values (t = 0). Error bars represent the standard deviation. Statistical significance was assessed by Student’s t-test. **C)** SDS PAGE and Western blot analysis with the indicated antibodies of whole cell protein extracts from the indicated strains expressing *ART2-HTF*. Cells were starved (- N) for 3h after 24h exponential growth. **D)** Live-cell fluorescence microscopy analysis of the indicated strains expressing *MUP1-GFP* from plasmid. Cells were starved (- N) for 6h after 24h exponential growth. **E), F)** The indicated strains were analyzed as in C) (upper panels) and D) (lower panels). **G)** Live-cell fluorescence microscopy analysis of *art2Δ* or *gcn4Δ* cells expressing pRS415-*MUP1-GFP* and pRS416-*pTDH3-ART2* starved (- N) for 6h after 24h exponential growth. Scale bars = 5 µm. See also Fig. S4.

In search of the pathway responsible for induction of Art2, we queried the results of our screen and identified the eIF2 kinase Gcn2 and several more *GCN* genes encoding key components of the general amino acid control (GAAC) pathway (Table S2). Gcn2 is activated during amino acid starvation by unloaded tRNAs and subsequently phosphorylates eIF2α to reduce protein synthesis in general, but thereby stimulates specifically the translation of the transcription factor Gcn4. Gcn4 then activates the transcription of genes, many of which are involved in amino acid metabolism (Hinnebusch, 2005). *gcn4Δ* was not scored in our genome-wide screen because it failed the quality control due to its slow growth phenotype. Interestingly, the promoter of *ART2* contained predicted Gcn4 binding sites (Venters et al., 2011, Schuldiner et al., 1998). Disrupting the GAAC pathway (*gcn4Δ*) eliminated the induction of Art2 in response to starvation at mRNA and protein levels (Fig. 4B,C). Consistently, the starvation-induced endocytosis of Mup1-GFP was hampered in *gcn4Δ* cells and several other *gcn* mutants (Fig. 4D, S4B). Also, starvation-induced endocytosis of Can1 was dependent on the GAAC pathway (Fig. S4C). When we introduced mutations in the predicted Gcn4 binding sites in the *ART2* promoter, Art2 protein levels no longer increased in response to starvation, and starvation-induced endocytosis of Mup1-GFP was impaired (Fig. 4E).

The expression of a constitutively translated Gcn4^C^ construct (Mueller and Hinnebusch, 1986) increased Art2 protein levels already under rich conditions, as revealed by WB analysis (Fig. 4F, compare Art2 protein levels in lanes 1 and 3), and drove unscheduled Mup1-GFP endocytosis (Fig. 4F, lower panel). Consistently, over-expression of Art2 in WT cells or in *gcn4Δ* mutants using the strong and constitutively active *TDH3* promoter (Fig. S4D) initiated Mup1-GFP endocytosis already under nutrient replete conditions (Fig. 4G) and bypassed the requirement of Gcn4 during starvation.

These results indicated that during amino acid starvation, Art2 transcription was induced via the GAAC pathway, leading to the up-regulation of Art2 protein levels. Moreover, it seemed that the up-regulation of Art2 protein levels was sufficient to drive Mup1 endocytosis. Hence, under amino acid replete conditions Art2 must be tightly repressed to prevent AAT endocytosis.

### Art2 directs Rsp5-dependent ubiquitination of C-terminal lysine residues in Mup1

To determine how Art2 contributed to Mup1 endocytosis, we examined its role in Mup1 ubiquitination. WT cells were harvested and Mup1-GFP was immunoprecipitated in denaturing conditions before and at different time points after starvation. Equal amounts of immunoprecipitated full-length Mup1-GFP were subjected to SDS-PAGE and WB analysis to compare the extent of its ubiquitination at different time points (Fig 5A). This analysis indicated that a pool of Mup1 was ubiquitinated prior to the onset of starvation (Fig. 5A, lane 1). At the onset of starvation, ubiquitination of Mup1-GFP appeared to decrease for some time (Fig. 5A lanes 2-4), until ubiquitination of Mup1-GFP began to increase again after 2-3 hours during starvation (Fig. 5A lanes 4-6), temporally coinciding with Art2 induction and starvation-induced endocytosis. Mup1-GFP was still ubiquitinated in *art2Δ* cells growing under rich conditions (Fig. 5B lane 4) and seemingly de-ubiquitinated at the onset of starvation, but the increase of ubiquitination during starvation was no longer observed (Fig. 5B, lane 6). Hence, Art2 was essential for the starvation-induced ubiquitination of Mup1.

**Figure 5:**
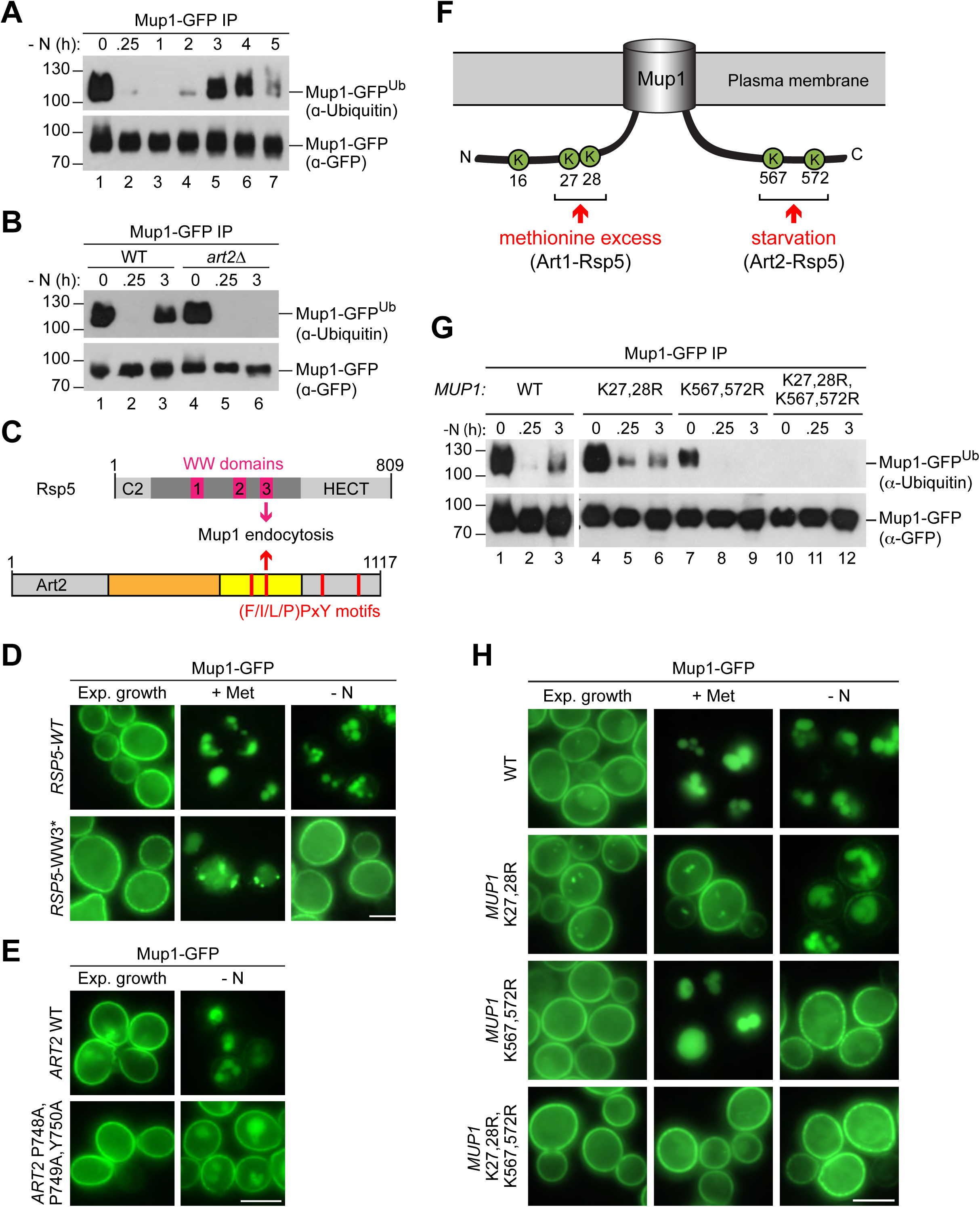
Art2-Rsp5 mediates the starvation-induced ubiquitination of Mup1-GFP at two specific C-terminal lysine residues. **A), B)** SDS PAGE and Western blot analysis with the indicated antibodies of immunoprecipitated Mup1-GFP from WT cells or *art2Δ* cells starved for the indicated times after 24h of exponential growth. Equal amounts of immunoprecipitated Mup1-GFP were loaded to compare the extent of ubiquitination. **C)** Scheme depicting the domain arrangement of Rsp5 and Art2, indicating the localization of the WW domains and PY motifs required for starvation-induced endocytosis of Mup1. **D)** Live-cell fluorescence microscopy analysis of *rsp5*Δ cells expressing pRS416-*MUP1-GFP* and pRS415- *HTF-RSP5-WT* (wild type) or pRS415-*HTF-RSP5-WW3\**. Cells were treated with 20 µg/ml L-methionine (+ Met) for 1.5h or starved (- N) for 6h after 24h exponential growth. **E)** Live-cell fluorescence microscopy analysis of *art2*Δ cells expressing *MUP1-GFP* and pRS416-*ART2* (WT) or pRS416-*ART2 P748A,P749A,Y750A*. Cells were starved (- N) for 6h after 24h exponential growth. **F)** Scheme of Mup1 topology with the N- and C-terminal ubiquitination sites targeted during substrate excess by Art1-Rsp5 and during starvation by Art2-Rsp5. Ubiquitinated lysines (K) shown in green with numbers corresponding to amino acid positions in the Mup1 sequence. **G)** WT cells expressing *MUP1-GFP* WT or the indicated *MUP1-GFP* mutants starved for the indicated times after 24h of exponential growth analyzed as in B). **H)** Live-cell fluorescence microscopy analysis of cells expressing *MUP1-GFP* (wild type (WT)), *MUP1 K27,28R, MUP1 K567,572R-GFP or MUP1 K27,28R,567,572R-GFP* as in D). Scale bars = 5 µm. See also Fig. S5.

α*−*Arrestins use PY motifs to bind to at least one of the three WW domains of Rsp5 (Lin et al., 2008). Starvation-induced endocytosis of Mup1 (but not methionine-induced endocytosis) was particularly dependent on the WW3 domain of Rsp5 (Fig. 5C, D, S5A). Art2 has four putative PY motifs (Fig. 5C) and the PY motif within the predicted arrestin fold of Art2 (P748,P749,Y750) was required for starvation-induced endocytosis (Fig. 5E). The Art2^P748A,P749A,Y750A^ mutant was expressed at similar levels than the WT protein (Fig. S5B). We suggest that interaction between WW3 in Rsp5 and the PY motif (748-750) of Art2 was required for the starvation-induced endocytosis of Mup1.

To identify lysine residues in Mup1 that were ubiquitinated in response to starvation, Mup1-GFP was immunoprecipitated before and three hours after starvation, digested and subjected to liquid chromatography-mass-spectrometry (LC-MS) (Fig S5C). Two N-terminal lysine residues (K16 and K27) and two C-terminal lysine residues (K567 and K572) in Mup1 were found to be ubiquitinated (Fig. 5F. Fig S5C), consistent with available high-throughput proteomic datasets (Swaney et al., 2013, Iesmantavicius et al., 2014). Earlier reports showed that two N-terminal lysine residues (K27, K28) in Mup1 were ubiquitinated by Art1-Rsp5 and required for methionine-induced endocytosis (Guiney et al., 2016, Busto et al., 2018, Ghaddar et al., 2014) (Fig. 5F). The role of ubiquitination at C-terminal lysine residues was not clear.

To determine how these N- and C-terminal lysine residues contributed to Mup1 ubiquitination under different growth conditions, we mutated the N-terminal (K27, K28) or C-terminal (K567, K572) lysine residues to arginine and additionally generated a quadruple mutant (K27,28,567,572R). Equal amounts of immunoprecipitated Mup1-GFP, Mup1^K27,28R^-GFP, Mup1^K567,572R^-GFP and Mup1^K27,28,567,572R^-GFP from exponentially growing cells or after starvation or methionine treatment were subjected to SDS-PAGE and WB analysis to compare their ubiquitination (Fig. 5G, S5D). Preventing the ubiquitination of the C-terminal lysine residues (K567,572R) abrogated starvation-induced ubiquitination of Mup1 (Fig. 5G, lanes 7-9), whereas Mup1^K27,28R^-GFP was still ubiquitinated after starvation (Fig. 5G, lanes 4-6). Upon methionine treatment, Mup1^K567,572R^-GFP ubiquitination was comparable to WT Mup1 (Fig. S5D, lanes 1-4), while the ubiquitination of Mup1^K27,28R^ was impaired as reported previously (Fig. S5D, lanes 5,6) (Guiney et al., 2016, Busto et al., 2018, Ghaddar et al., 2014). The quadruple mutant Mup1^K27, 28, 567, 572R^-GFP always appeared to be devoid of ubiquitination (Fig. 5G, lanes 11,12; Fig. S5D, lanes 7,8), suggesting that the two N-terminal and the two C-terminal lysine residues are the major sites for ubiquitination. Consistently, the quadruple lysine mutation rendered Mup1^K27,28,567,572R^-GFP refractory to methionine- and starvation-induced endocytosis (Fig. 5H). Mutations in the N-terminal lysine residues (K27,28R) blocked methionine-induced endocytosis, but not starvation-induced endocytosis. In contrast, starvation-induced endocytosis of Mup1^K567,572R^-GFP was specifically blocked, but not its methionine-induced endocytosis (Fig. 5H).

Collectively, these results suggested that Art2 directs Rsp5 to the ubiquitination of the C-terminal lysine residues K567 and K572 during starvation, whereas Art1-Rsp5 complexes ubiquitinated Mup1 on the two N-terminal lysine residues K27 and K28 in response to methionine excess (Fig. 5F).

### The C-terminus of Mup1 contains an acidic sorting signal for Art2

In addition to ubiquitination, our mass-spectrometry analysis of immunoprecipitated Mup1-GFP from starved cells identified potential phosphorylation sites at the N-terminus (S9,23,31,42 and T34) and at the C-terminus of Mup1 (T552 and S568,573) (Fig. 6A, S5C). Additional phosphorylation on T560 was reported in other studies (Swaney et al., 2013, Iesmantavicius et al., 2014). Of note, the C-terminal phosphorylation sites were proximal (T552, T560) or directly adjacent (S568, S573) to lysine residues K567 and K572 that were ubiquitinated in an Art2-dependent manner in response to starvation (Fig. 6A, S5C).

**Figure 6:**
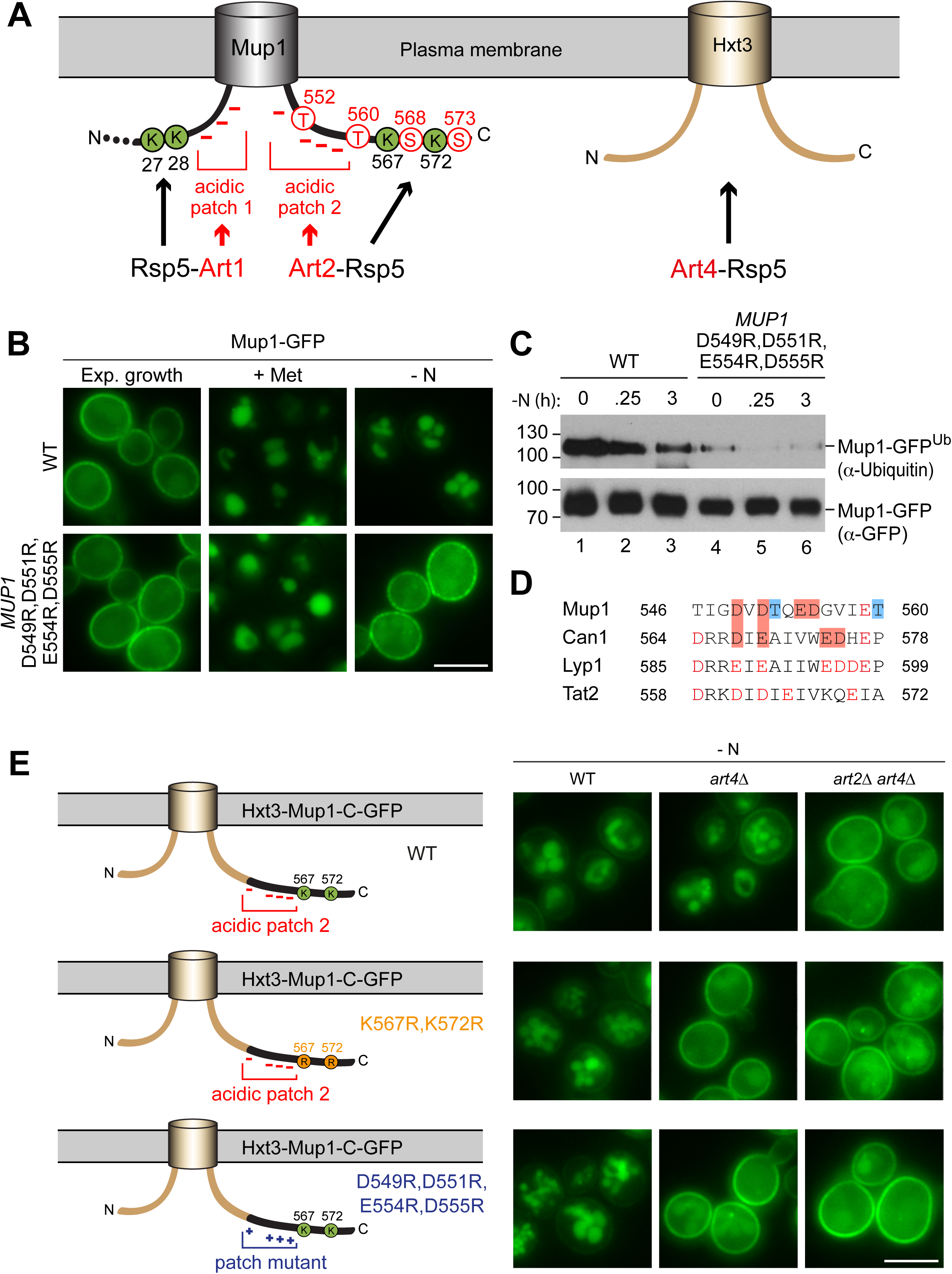
The C-terminus of Mup1 harbors a transplantable, starvation-responsive acidic degron. **A)** Left: scheme of Mup1 topology with N- and C-terminal ubiquitination sites and acidic patches targeted by Art1-Rsp5 and Art2-Rsp5, respectively, and the C-terminal phosphorylation sites of Mup1 that promote its starvation-induced endocytosis. Ubiquitinated lysines (K) shown in green and phosphorylated serines (S) and threonines (T) in red with numbers corresponding to amino acid positions in the Mup1 sequence. Right: Hxt3 as an Art4-Rsp5 dependent cargo during nitrogen starvation. **B)** Live-cell fluorescence microscopy analysis of Mup1-GFP endocytosis in cells expressing *MUP1-GFP* (wild type (WT)) or *MUP1 D549R,D551R,E554R,D555R-GFP*. Cells were treated with 20 µg/ml L-methionine (+ Met) for 1.5h or starved (- N) for 6h after 24h exponential growth. **C)** SDS PAGE and Western blot analysis with the indicated antibodies of immunoprecipitated Mup1-GFP from cells expressing *MUP1-GFP* (WT) or *MUP1 D549R,D551R,E554R,D555R -GFP* starved for the indicated times after 24h of exponential growth. Equal amounts of immunoprecipitated Mup1-GFP were loaded to compare the extent of ubiquitination. **D)** Amino acid sequence alignment of the C-terminal acidic patches of Mup1, Can1, Lyp1 and Tat2. The boxes indicate acidic residues (red) and phosphorylation sites (blue), which are required for Art2-dependent starvation-induced endocytosis. Red letters illustrate further acidic residues. **E)** Live-cell fluorescence microscopy analysis of wild type (WT), *art4*Δ and *art2*Δ *art4*Δ cells expressing *HXT3-MUP1-C-GFP* (top panels), *HXT3-MUP1-C K567,572R-GFP* (middle panels) or *HXT3-MUP1-C D549R,D551R,E554R,D555R -GFP* (bottom panels). Cells were starved (- N) for 6h after 24h exponential growth. Scale bars = 5 µm. See also Fig. S6.

Mutation of the C-terminal threonine residues to alanine (T552,560A) to prevent phosphorylation specifically blocked starvation-induced endocytosis of Mup1^T552,560A^-GFP, but not methionine-induced endocytosis (Fig. S5E). Mutation of these phosphorylation sites also prevented ubiquitination of Mup1 during starvation (Fig. S5F). Blocking the phosphorylation of S568 and S573 also impaired starvation-induced endocytosis of Mup1^S568,573A^-GFP (Fig. S5E), yet starvation-induced ubiquitination could still be detected (Fig. S5F). Mutation of most N-terminal serine and threonine residues, including all experimentally determined phosphorylation sites, to alanine (T6,25,26,34A,S9,22,23,24,31,33,42A) did not affect starvation-induced endocytosis of Mup1 (Fig. S5G). It seemed that the phosphorylation of C-terminal threonine residues (T552, 560) contributed to the starvation-induced Art2-Rsp5 ubiquitination of Mup1 and thus might add an additional layer of regulation, similar to the recognition of GPCRs by β–arrestins in mammalian cells (Shukla et al., 2014, Nikko et al., 2008, Marchal et al., 2000, Hicke et al., 1998).

An extended acidic Art1 sorting motif at the N-terminus of Mup1 is required for ubiquitination of K27,28 and subsequent Mup1 endocytosis in response to methionine excess (Guiney et al., 2016, Busto et al., 2018). Consistent with our model involving two distinct mechanisms for Mup1 endocytosis, mutations in the acidic N-terminal Art1 sorting motif of Mup1 (D43A,G47A,Q49A,T52A,L54A) blocked Art1-Rsp5-dependent methionine-induced endocytosis, but not Art2-Rsp5-dependent starvation-induced endocytosis (Fig. S6A).

The C-terminus of Mup1 also contains an acidic patch (D549-D555), close to the ubiquitination sites K567 and K572 and the C-terminal threonine phosphorylation sites (T552, 560) involved in starvation-induced endocytosis (Fig. 6A, B). Mutation of the acidic residues in this region to basic amino acids (R) demonstrated that this C-terminal acidic region was specifically required for starvation-induced endocytosis. Live cell fluorescence microscopy revealed that Mup1^D549R,D551R,E554R,D555R^-GFP remained at the PM in response to starvation, whereas methionine-induced endocytosis was not impaired (Fig. 6B). Even Art2 overexpression failed to induce endocytosis during exponential growth or starvation when the C-terminal acidic patch in Mup1 was mutated (Fig. S6B). Moreover, immunoprecipitation of Mup1^D549R,D551R,E554R,D555R^-GFP and subsequent SDS-PAGE and WB analysis revealed that it was no longer efficiently ubiquitinated (Fig. 6C, lanes 4 - 6), suggesting that the C-terminal acidic patch was essential for the Art2-Rsp5-dependent ubiquitination during starvation.

Comparing the amino acid sequences of the C-terminal tails of the four Art2-dependent cargoes, Mup1, Can1, Tat2 and Lyp1, indicated similar acidic patches (Fig. 6D). To analyze if the acidic patch in Can1 also contributed to starvation-induced endocytosis, we mutated D567, E569, E574 and E575 to arginine. Mutant Can1^D567R,E569R,E574R,D575R^-GFP mostly localized to the PM under growing conditions. Importantly, the Art2-dependent starvation-induced down-regulation of Can1^D567R,E569R,E574R,D575R^-GFP was impaired (Fig. S6C). These results imply that Mup1, Can1 and potentially also Lyp1 and Tat2 have acidic amino acid sequences at their C-termini that could serve as sorting signal for Art2-mediated starvation-induced endocytosis.

### The C-terminal acidic sorting signal of Mup1 is sufficient for Art2-dependent starvation-induced endocytosis

It seemed that the last 26 amino acid residues (aa 549–574) of Mup1 harbor three features that are collectively required specifically for starvation-induced endocytosis: putative phosphorylation sites, the acidic patch and the ubiquitination sites. Hence, we tested if the C-terminal region of Mup1 was sufficient to convert an Art2-independent cargo into an Art2 cargo. We selected the low affinity glucose transporter Hxt3, which was efficiently removed from the PM in response to starvation (Fig. S6D, Fig.1B, Table S1). Live cell fluorescence microscopy and WB analysis showed that starvation-induced endocytosis of Hxt3-GFP was independent of Art2, but instead required Art4 (Fig. S6D). In *art4Δ* mutants, but not in *art2Δ* mutants, Hxt3-GFP remained mostly at the PM (Fig. S6D) in response to starvation, and its vacuolar degradation was impaired (Fig. S6E, lane 4). However, when the C-terminal 30 amino acids of Mup1 (aa 545-574) were fused onto the C-terminus of Hxt3 (Fig. 6E), they restored starvation-induced endocytosis in *art4Δ* mutants. This Hxt3-Mup1-C-GFP chimera was now efficiently removed from the PM and transported to the vacuole in the vast majority of *art4Δ* cells (86%, n=76 cells) (Fig. 6E), most likely because it became an Art2 substrate. Indeed, the additional deletion of *ART2* (*art2Δ art4Δ)* blocked starvation-induced endocytosis and demonstrated that Art2 now mediated the degradation of Hxt3-Mup1-C-GFP in *art4Δ* mutants. The degradation of Hxt3-Mup1-C-GFP and the concomitant accumulation of free GFP were further examined by SDS-PAGE and WB analysis from total cell extracts (Fig. S6F). This analysis showed that the majority of the full length Hxt3-Mup1-C-GFP chimera was degraded in WT cells, *art2Δ* and *art4Δ* mutants during starvation (Fig. S6F, lanes 2, 4, 8), but only poorly in *art2Δ art4Δ* double mutants (Fig. S6B, lane 6).

The Art2-dependent endocytosis of the Hxt3-Mup1-C-GFP chimera required two key features provided by the C-terminus of Mup1 (the acidic patch and the two C-terminal lysine residues), since in *art4Δ* cells starvation-induced endocytosis of Hxt3-Mup1-C^K567,572R^-GFP and Hxt3-Mup1-C^D549R,D551R,E554R,D555R^-GFP was blocked (Fig. 6E).

Taken together, these results demonstrate that the C-terminus of Mup1 (aa 545–574) encodes a portable acidic sorting signal that can be recognized by Art2 and directs Rsp5 to ubiquitinate specifically two proximal lysine residues to promote starvation-induced endocytosis.

### A basic patch of Art2 is required for starvation-induced degradation of Mup1

After having defined that the C-terminus of Mup1 (and possibly also the C-terminus of the AATs Can1, Lyp1 and Tat2) provides a degron sequence for Art2-Rsp5 complexes, we addressed how it could be specifically recognized. Upon inspection of the predicted arrestin domain in Art2, we noted a stretch of positively charged residues within the arrestin-C domain (Fig. 7A). Converting these basic residues into an acidic patch (Art2^K664D,R665D,R666D,K667D^) abolished starvation-induced endocytosis of Mup1 (Fig. 7B). Western blot analysis of total cell lysates showed that the Art2 basic patch mutant protein was expressed at similar levels as WT Art2 and was also upregulated after 3 hours of starvation (Fig. S7A, lane 6). In addition, the Art2 basic patch mutant also impaired, at least partially, starvation-induced endocytosis of Can1 and Lyp1, while the endocytosis of Tat2 was independent of the basic patch (Fig. 7B, S7B).

**Figure 7:**
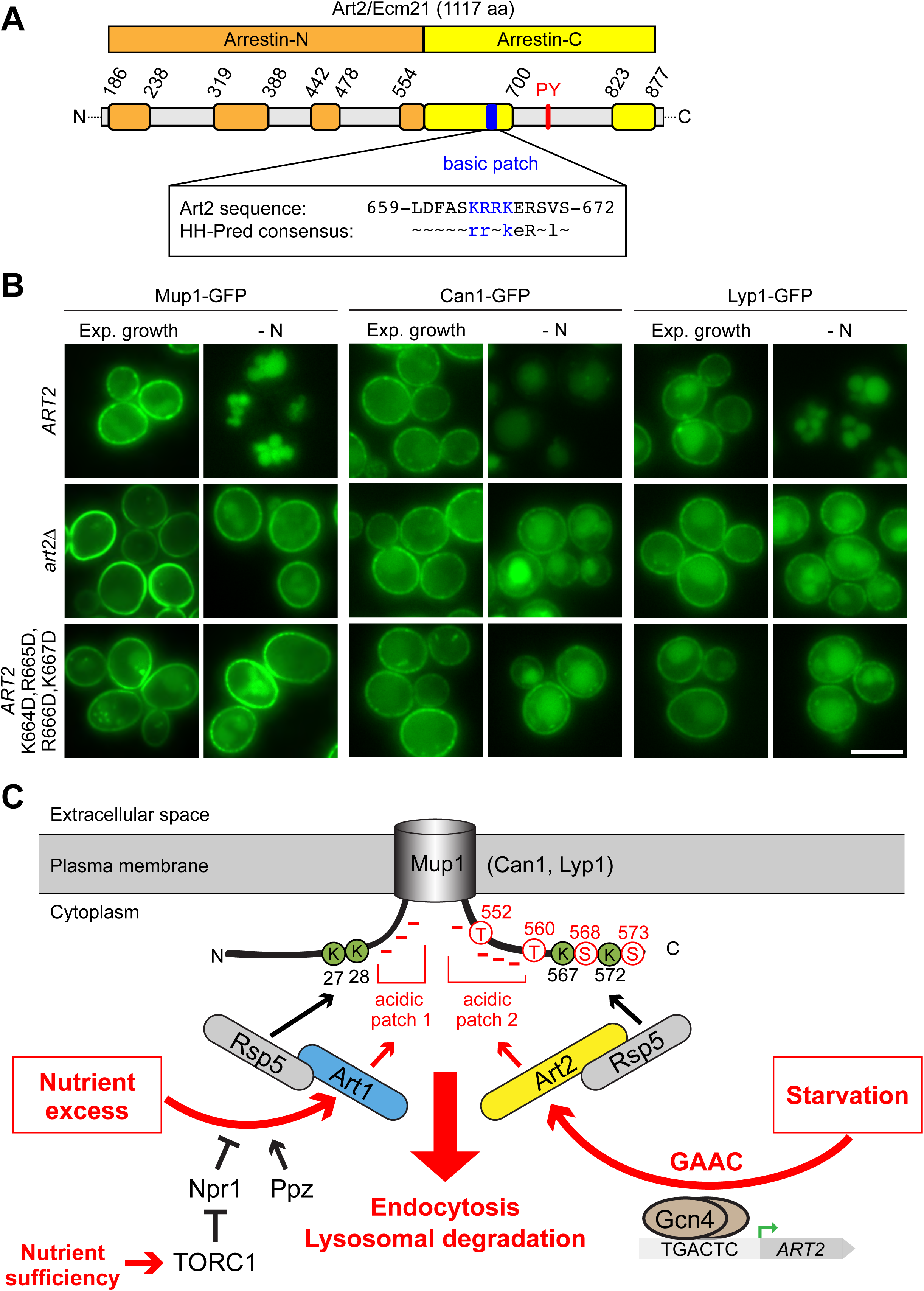
A basic patch of Art2 promotes the starvation-induced endocytosis of Mup1, Can1 and Lyp1. **A)** Scheme of Art2 topology with arrestin-N domain in orange, arrestin-C domain in yellow and tails and interspersed extended loops (Baile et al., 2019) in light grey. The basic amino acid residues shown in blue mediate the starvation-induced endocytosis of Mup1, Can1 and Lyp1 (numbers correspond to amino acid positions in the Art2 sequence). Below is aligned the HHpred consensus sequence for Art2/Ecm21 derived from three HHblits iterations (Zimmermann et al., 2018), suggesting conservation of the basic patch in the Art2 protein family. **B)** Live-cell fluorescence microscopy analysis of *art2*Δ cells expressing *MUP1-GFP*, *CAN1-GFP* or *LYP1-GFP* and pRS416-*ART2-WT*, empty vector or *pRS416-ART2 K664D,R665D,R666D,K667D*. Cells were starved (- N) for 6h after 24h exponential growth. **C)** Scheme for the regulation of the substrate- and starvation-induced endocytosis of Mup1. During substrate excess TORC1 inhibits the Npr1 kinase which otherwise would phosphorylate and inhibit Art1. Art1-Rsp5 becomes dephosphorylated and subsequently binds the acidic patch 1 at the N-terminal tail of Mup1, leading to the ubiquitination of K27 and K28 and degradation of Mup1 (left). During amino acid and nitrogen starvation the general amino acid control (GAAC) pathway upregulates the ubiquitin ligase adaptor Art2 via the transcriptional regulator Gcn4. The ensuing Art2-Rsp5 complex binds with its basic patch to the acidic patch 2 of Mup1 leading to the ubiquitination of K567 and K572 and degradation of Mup1 (right). At the same time, starvation causes TORC1 inhibition and activation of Npr1 and inhibits Art1-dependent ubiquitination. Ubiquitinated lysines (K) shown in green and phosphorylated serines (S) and threonines (T) in red with numbers corresponding to amino acid positions in the Mup1 sequence. Scale bars = 5 µm. See also Fig. S7.

Overall, it seemed that Art2 employed a positively charged region in its arrestin fold to recognize C-terminal acidic patches in at least three different AATs (Mup1, Can1, Lyp1) and thus mediate their endocytosis in starvation conditions.

## Discussion

We have made progress towards understanding how yeast cells selectively re-configure their repertoire of transporters at the PM in response to their nutritional status. The model in Fig. 7C provides the conceptual framework for the mutually exclusive but complementary action of Art1-Rsp5 and Art2-Rsp5 ubiquitin ligase complexes for mediating the selective endocytosis of the methionine transporter Mup1 in response to changes in amino acid availability. Similar concepts may apply for the arginine transporter Can1, the lysine transporter Lyp1 and the tryptophan and tyrosine transporter Tat2. Based on our results and work from others, the endocytic down-regulation of these AATs can be described by the following set of rules: (1) Reciprocal (de-)activation of the α*−*arrestins Art1 and Art2. (2) Art1 or Art2 each recognize different acidic sorting signals on their client AATs that may require additional phosphorylation. (3) To do so, they employ basic patches in their extended arrestin C-domains and defined PY motifs to orient Rsp5 with high specificity towards proximal lysine residues. These rules satisfy the plasticity required for different α*−*arrestin and AAT interactions that drive exclusive or relatively broad substrate specificity depending on the metabolic context.

While both Art1 and Art2 lead to the degradation of AATs, they answer to distinct metabolic cues and are thus wired into distinct signaling pathways. Activation of Art1 by amino acid influx requires the coordinated interplay of TORC1 signaling to inactivate Npr1 (a kinase that negatively regulates Art1) and the action of phosphatases (Gournas et al., 2017, Lee et al., 2019, MacGurn et al., 2011, Tumolo et al., 2020). In response to amino acid limitation, TORC1 is no longer active. This will activate Npr1 to phosphorylate Art1, thereby inactivating it. At the same time, the lack of amino acids will activate the eIF2α kinase Gcn2. Gcn2 will phosphorylate eIF2α, which leads to the global down-regulation of translation, but enables specific translation of the transcription factor Gcn4 (Hinnebusch, 2005). Gcn4 then induces transcription of genes required for amino acid biosynthesis and of *ART2*, which causes an increase in Art2 protein levels and thus formation of Art2-Rsp5 complexes. This appears as primary means to activate Art2, since unscheduled increase in Art2 protein levels was sufficient to drive Art2-dependent nutrient transporter endocytosis already in cells growing under rich conditions. When amino acids become available again, the system can efficiently reset. TORC1 is reactivated resulting in Art1 reactivation. Conversely, Gcn4 will become instable and rapidly degraded by the UPS (Kornitzer et al., 1994, Meimoun et al., 2000, Irniger and Braus, 2003), and thus, the transcription of *ART2* will cease. Interestingly, two de-ubiquitinating enzymes (Ubp2, Ubp15) de-ubiquitinate Art2 to influence its protein stability (Ho et al., 2017, Kee et al., 2006). Inhibiting their activity could provide additional control to repress Art2-dependent endocytosis in cells growing under rich conditions. Our screen identified also two de-ubiquinating enzymes, Doa4 and Ubp6 to be specifically required for starvation-induced endocytosis of Mup1. They could act directly on Art2 or Mup1 or help to maintain homeostasis of the ubiquitin pool during starvation.

Art2 is subject to extensive post-translational modification, including ubiquitination and phosphorylation. Database searches and our own proteomic experiments identified 68 phosphorylation sites and 20 ubiquitination sites in Art2 (data not shown) (Swaney et al., 2013, Albuquerque et al., 2008, Holt et al., 2009). How these modifications help to control the activity of Art2 remains a complex and open questions. Several arrestins were found to be phospho-inhibited in specific conditions (MacGurn et al., 2011, Becuwe et al., 2012b, O’Donnell et al., 2013, Hovsepian et al., 2017, Merhi and André, 2012, Llopis-Torregrosa et al., 2016), the common molecular basis of which is unknown. An exciting hypothesis would be that α-arrestin hyper-phosphorylation would add negative charges, and thereby prevent the recognition of acidic patches on transporters through electrostatic repulsion. Interestingly, our screen identified the pleitropic type 2A-related serine-threonine phosphatase Sit4 as a class 1 hit. Hence, Sit4 may be linked directly or indirectly to de-phosphorylation of Art2 and controlling its activity as reported recently for the Art2-dependent regulation of vitamin B1 transporters (Savocco et al., 2019).

Through the complementary activation of Art1 and Art2 cells can coordinate amino acid uptake through at least four high-affinity amino acid transporters with amino acid availability. The regulation of hexose transporters by glucose availability appears to be conceptually related, with distinct α*−*arrestin-Rsp5 complexes in charge of down-regulating the same transporters at various glucose concentrations with distinct mechanisms and kinetics (Hovsepian et al., 2017, Nikko and Pelham, 2009). In particular, the endocytosis of high-affinity hexose transporters during glucose starvation involves Art8, the closest paralogue of Art2, whose expression is also controlled by nutrient-regulated transcription (Hovsepian et al., 2017). Altogether, a picture emerges in which the transcriptional control of α*−*arrestin expression by nutrient-signaling pathways is critical to cope with nutrient depletion.

Our work also extends on previous findings regarding the determinants of α*−*arrestin/transporter interaction, indicating communalities between starvation- and substrate-induced endocytosis. Art1-Rsp5 and Art2-Rsp5 complexes both recognize specific acidic sequences on Mup1 (Fig. 7C). Activated Art1-Rsp5 complexes recognize an acidic stretch in the N-terminus of Mup1 and Can1, close to the first transmembrane domain. The exposure of these N-terminal acidic patches is linked to substrate transport and conformational transitions in the transporters from the outward-open to the inward-open conformation, which in Mup1 also includes the so-called ‘C-plug’ (aa 520–543) (Busto et al., 2018, Guiney et al., 2016, Gournas et al., 2017). This conformational switch drives lateral re-localization of Mup1 and Can1 into a disperse PM compartment, where they are ubiquitinated by Art1-Rsp5 (Gournas et al., 2018, Busto et al., 2018). Art2 recognizes specifically an acidic patch in the C-terminal tail of Mup1, and thereby directs Rsp5 to ubiquitinate two juxtaposed C-terminal lysine residues. The C-plug is very close to the C-terminal acidic patch, but is not part of the C-terminal Mup1 degron. We speculate that in the absence of nutrients AATs will spend more time in the outward open state with the C-Plug in place. In this state, activated Art2-Rsp5 complexes can still engage the C-terminal acidic patches. Hence, toggling Art1/Art2 activation in combination with accessibility of N- or C-terminal acidic sorting signals in AATs, in part regulated by their conformational state, must fall together to allow selective endocytosis.

An additional layer of regulation for endocytosis is provided by phosphorylation of AATs close to the acidic sorting signal. At the moment we can only speculate about the kinase responsible for the phosphorylation of the C-terminal serine or threonine sites of Mup1. Perhaps, constitutive PM-associated kinases such as the yeast casein kinase 1 pair (Yck1/2) are involved, which are known to recognize rather acidic target sequences and to regulate endocytosis (Hicke et al., 1998, Paiva et al., 2009, Nikko et al., 2008, Marchal et al., 2002).

α-Arrestins lack the polar core in the arrestin domain that is used for cargo interactions in β-arrestins (Aubry et al., 2009, Polekhina et al., 2013). Instead Art1-Rsp5 and Art2-Rsp5 complexes each use a basic region in their arrestin C-domain to detect the acidic sorting signal in their client AATs. Studies on the interaction between GPCRs and β-arrestins revealed a multimodal network of flexible interactions: The N-domain of β-arrestin interacts with phosphorylated regions of the GPCR, their finger loop inserts into the transmembrane domain bundle of the GPCR and loops at the C-terminal edge of β-arrestin engage the membrane (Staus et al., 2020, Huang et al., 2020). Perhaps a similar concept also holds true for α-arrestins. This is not unlikely given that their arrestin fold appears to be interspersed with disordered loops and very long, probably unstructured N- and/or C-terminal tails, some of which participate in cargo recognition or membrane interactions (Baile et al., 2019).

Despite the possible plasticity in substrate interactions, the selectivity of Art1-Rsp5 and Art2-Rsp5 complexes in ubiquitinating lysine residues proximal to the acidic patches of Mup1 is remarkable. Mup1 has 19 lysine residues at the cytoplasmic side: four at the N-terminal tail, six in the C-terminal tail and 9 in the intracellular loops of the pore domain. Yet, Art1-Rsp5 complexes only ubiquitinate K27 and K28, whereas Art2-Rsp5 complexes only ubiquitinate K567 and K572. Also in the Hxt3-Mup1-C chimeric protein, Art2-Rsp5 complexes ubiquitinated only the lysine residues close to the acidic patch, despite 6 further lysine residues in the directly adjacent C-terminal tail of Hxt3. How is this possible? We speculate that these two α-arrestin-Rsp5 complexes orient the HECT domain of Rsp5 with high precision towards the lysine residues that are spatially close to the acidic patches. Once ubiquitinated, the AAT can engage the endocytic machinery to be removed from the PM.

In conclusion, Art1-Rsp5 complexes act rapidly to prevent the accumulation of excess amino acids, whereas the Art2-Rsp5 complexes help to degrade idle high affinity amino acid transporters over longer periods of starvation to recycle their amino acid content. Starvation-induced endocytosis and the subsequent degradation of membrane proteins is required to maintain intracellular amino acid homeostasis (Müller et al., 2015, Jones et al., 2012). As such, it is well suited that Art2 activity and thus starvation-induced endocytosis is co-regulated and coordinated with *de novo* amino acid biosynthesis via the GAAC pathway. The down-regulation of AATs together with glucose transporters and further PM proteins could also free up domains at the PM that are populated by selective nutrient transporters (Spira et al., 2012, Grossmann et al., 2008) for transporters with broader substrate specificity such as the general amino acid permease Gap1 and the ammonium transporter Mep2, which are strongly up-regulated during starvation. Hence starvation-induced endocytosis could prepare cells – anticipatory – for non-selective nutrient acquisition, as soon as nutrients become available again.

Altogether, our results provide a better understanding of the molecular mechanisms that couple metabolic signaling and nutrient availability to nutrient transporter endocytosis in budding yeast. This might provide basic insights and principles for the vastly more complex regulation of the nutrient transporter repertoire in human cells under normal as well as pathophysiological conditions.

## Material and Methods

### Yeast strains, media and growth conditions

Yeast strains used for the microscopy screen for starvation-responsive endocytosis cargoes were mainly derived from the Yeast C-terminal GFP Collection (Huh et al., 2003) with addition of further C-terminally-tagged transporters in BY4741 (*MATa his3Δ1 leu2Δ0 met15Δ0 ura3Δ0*) and SEY6210 (*MATα leu2-3, 112 ura3-52 his3-200 trp1-901 lys2-801 suc2-9*) strain background (table S1). The FACS screen for genes affecting the starvation-induced endocytosis of Mup1-pHluorin was performed using the non-essential gene deletion strain collection purchased from Open Biosystems (*BY4742: MATα his3Δ1 leu2Δ0 lys2Δ0 ura3Δ0*) transformed with pRS416 expressing Mup1-pHluorin. For all other experiments the SEY6210 parental strain was used (table S4).

In all experiments except the genome-wide FACS screen cells were cultivated in rich medium containing: 6,7g/L Yeast Nitrogen Base without amino acids (#HP26.1, Roth, Germany), 20mg/L adenine hemisulfate (#A3159-25G, Sigma, Austria), 20mg/L arginine (#1655.1, Roth, Germany), 230 mg/L lysine monohydrate (#4207.1, Roth, Germany), 300 mg/L threonine (#T206.2, Roth, Germany), 30 mg/L tyrosine (#T207.1, Roth, Germany), 2% glucose (#X997.5, Roth, Germany). The rich medium was supplemented when required for the auxotrophic strains with: 20 mg/L histidine (#3852.1, Roth, Germany), 60 mg/L leucine (#3984.1, Roth, Germany), 20 mg/L tryptophan (#4858.2, Roth, Germany), 20 mg/L uracil (#7288.2, Roth, Germany), 20 mg/L methionine (#9359.1, Roth, Germany). Starvation medium contained 1,7g/L Yeast Nitrogen Base without amino acids and ammonium sulfate (#J630-100G, VWR, USA) and 2% glucose (#X997.5, Roth, Germany).

Yeast cells were cultivated in rich medium and kept for at least 24h in exponential growth phase by dilution into fresh medium before onset of all experiments at mid-log phase. All cultivations were done at 26°C. Substrate-induced endocytosis of Mup1 was triggered by the addition of 20 mg/L methionine (#9359.1, Roth, Germany) and, unless otherwise stated, analyzed 1.5h after the treatment. For starvation experiments, exponentially growing cells (0.4-0.6 OD_600_/ml) were washed twice with and resuspended in starvation medium (1 OD_600nm_/ml) and incubated for the indicated times.

### Genetic modifications and cloning

Genetic modifications were performed by PCR and/or homologous recombination using standard techniques. All chromosomal tags were introduced at the C-terminus of the target ORFs to preserve the endogenous 5 prime regulatory sequences. Chromosomally modified yeast strains were analyzed by genotyping PCR and/or DNA-sequencing. Plasmid-expressed genes including their endogenous or heterologous promoters were amplified from yeast genomic DNA and cloned into centromeric vectors (pRS series). All plasmids were analyzed by DNA-sequencing. Standard techniques were used for yeast transformation, mating and tetrad analysis or haploid selection. Plasmids and primers are listed in table S4.

### Flow cytometry screen

The yeast non-essential knock out collection (YSC1054; Open Biosystems) was transformed in 96-well format using a standard lithium acetate/PEG-4000 yeast transformation protocol with a pRS416 plasmid encoding Mup1-pHluorin. Transformants were selected on agar plates, inoculated into 160 µl of rich medium (YNB medium lacking uracil as specified above containing 1.5x of all amino acids) and incubated for 14h shaking (180 rpm) at 26° in 96-well plates (#83.3924, Sarstedt, Germany). Each plate also contained the WT BY4742 parental strain and *art2Δ* as negative and positive controls. 60 µl of the overnight cultures were transferred to 96-deep-well plates (#951033405, 96/2000µl, white border, Eppendorf, Germany) containing 600 µl rich medium and further incubated for 5h. At this point 120 µl of culture referred to as ‘exponentially growing’ were transferred to standard 96-well plates and analyzed by flow cytometry. The remaining culture was harvested by centrifugation (1807 g; 3 min), the medium was aspirated, cells were washed twice with 600 µl starvation medium and recovered by centrifugation. Subsequently, cells were resuspended in 600 µl starvation medium and incubated shaking (180 rpm) for 18-22h. 200 µl nitrogen-starved culture were transferred to standard 96-well plates and analyzed by flow cytometry. All pipetting steps were performed using the MEA Robotic System (PhyNexus, USA). Flow cytometry was performed in 96-well format using the Guava easyCyte 8HT-System (Sr.No. 6735128143), EMD Millipore, Merck, Germany) with the following settings: Energy GRN: 72 plus (YEL=8; RED=8; NIR=8; RED2=8; NIR2=8), green channel, forward scatter FSC= 14; sideward scatter SSC= 28; 15.000-20.000 counts/sample, acquire: 40sec, 3sec mix prior to acquisition. GuavaSoft 2.7 software was used for data analysis. The positive/negative cut-off was set for each plate empirically at the intercept of the log/starvation histograms of the WT and *art2Δ* controls (*art2Δ*, which emerged as a well-reproducible hit early in the screen, was included as a negative control in all further plates). All potential hits were re-examined by fluorescence microscopy. To this end, at least 100 starved cells were analyzed by fluorescence microscopy after starvation and the percentage of cells showing a degradation-deficient phenotype (Mup1-pHluorin at the plasma membrane, in small cytosolic objects, class E-like objects or small objects within vacuoles) of the total number of cells counted was calculated (table S2). Strains with more than 45% cells with retained fluorescence after at least 18 hours of starvation were considered as hits. For a stringent final selection, we compared those hits to the original FACS screen and finally only considered those in which at least once more the 30% Mup1-pHluorin fluorescence was also retained after starvation in the FACS screen. In addition, most hits were also scored for methionine-induced endocytosis of Mup1-pHluorin. Hits were considered starvation-specific if the fluorescence was quenched in more than 67% of cells after 90 minutes of methionine treatment (20µg/ml).

### Gene ontology analysis

Gene Ontology (GO) enrichment analysis (Harris et al., 2004) was performed using 128 genes listed in Appendix Table S2. They were mapped against the generic GO-Slim: process, generic GO-Slim: component and macromolecular complex terms: component GO sets using the GoSLIM mapper of the *Saccharomyces* genome database. We calculated the ratio of the observed (cluster frequency) vs. the expected number of genes (genome frequency) associated with the GO term, referred to as enrichment over genome (McClellan et al., 2007). Only GO terms with more than one associated gene were reported. The full analysis is presented in Table S3.

### Fluorescence live cell wide-field microscopy

For microscopy, cells were concentrated by centrifugation and directly mounted onto glass slides. Live cell wide-field fluorescence microscopy was carried out using a Zeiss Axio Imager M1 equipped with a sola light engine LED light source (Lumencore), a 100x oil immersion objective (NA 1.45) standard GFP and mCherry fluorescent filters, a SPOT Xplorer CCD camera, and Visitron VisiView software (version 2.1.4). The brightness and contrast of the images in the figures were linearly adjusted using Photoshop CS5 (Adobe Version 12.0.4×64).

### Preparation of yeast whole cell protein extracts

10 OD_600nm_ yeast cells were pelleted by centrifugation, resuspended in ice-cold water with 10% trichloroacetic acid (#T0699, Sigma, Austria), incubated on ice for at least 30 min and washed twice with ice cold acetone (#5025.5, Roth, Germany). The precipitate was resuspended in 200 µl extraction buffer (150 mM Tris-HCl, pH 6,8 (#443866G, VWR, USA); 6% SDS (#CN30.3, Roth, Germany), 6M urea (#51456-500G, Sigma, Austria), 6% glycerol (#3783.2, Roth, Germany), 3% β-mercaptoethanol (#M6250, Sigma, Austria), 0,01% bromophenol blue (#44305, BDH Laboratory Supplies, England)) and solubilized with 0.75-1 mm glass beads (#A554.1, Roth Germany) for 15 min at RT and subsequent heating at 42°C for 30 min and 60°C for 10 min.

### Western blot and immunodetection

Protein extracts from total cells and eluates from immuno-precipitations were separated by standard SDS-PAGE and transferred to PVDF membranes (#10600023, VWR, USA) for the detection of Mup1-GFP, Art2-HTF and Pgk1 or to nitrocellulose membranes (#10600004, VWR, USA) for the detection of ubiquitinated Mup1. PVDF membranes were stained with Coomassie Brilliant Blue R250 (#3862.2, Roth, Germany) for assessment of transfer and loading. Antibodies used in this study include: α-FLAG M2 (#F1804, Sigma, Austria), α-GFP IgG1K (#11814460001, Sigma, Austria), α-Pgk1 22C5D8 (#459250, Invitrogen, USA), α-ubiquitin P4D1 (#3936S, Santa Cruz Biotechnology, USA).

### Immunoprecipitation

Mup1-GFP immunoprecipitation protocol was adapted from (Hovsepian et al., 2017). 50 OD_600nm_ yeast cells were harvested by centrifugation and mechanically disrupted by glass bead lysis (0.75-1 mm) at 4°C in 500 µl ice-cold RIPA lysis buffer (50 mM Tris-HCl, pH 7.5, 150 mM NaCl (#3957.5, Roth, Germany), 0.1% SDS, 2 mM EDTA (#ED-1KG, Sigma, Austria), 50 mM NaF (#SO0323, Scharlau, Spain), 1% Nonidet P 40 (#74385, Fluka, Germany), 0.5% Na-deoxycholate (#D6750-25G, Sigma, Austria), 1% glycerol) containing protease inhibitors (cOmplete Protease Inhibitor Cocktail (#11697498001, Sigma, Austria), yeast protease inhibitor cocktail (yPIC, #P8215-5ML, Sigma, Austria), 2 mM phenylmethylsulfonyl fluoride (PMSF, #P7626-5G, Sigma, Austria)) and 20 mM *N*-ethylmaleimide (NEM, #E3876-5G, Sigma, Austria) using four cycles of lysis (2 min), each separated by 2 min chilling on ice. The lysate was then rotated for 30 min at 4°C for solubilization. 500 µl wash buffer (50 mM Tris-HCl, pH 7.5, 150 mM NaCl, 1% Nonidet P 40, 5% glycerol) containing 20 mM NEM were added and the sample was mixed and further rotated for 1h at 4°C. After vortexing for 3 min, the lysate was centrifuged at 10000g for 10 min at 4°C. The cleared lysate was then added to 25 µl of GFP-Trap_MA magnetic agarose beads (#gtma-20, ChromoTek, Germany) prewashed twice in 1 ml wash buffer. The sample was rotated for 5h at 4°C. The beads were collected using a magnetic rack and washed twice by rotating for 15 min at 4°C with 1.5 ml ice-cold wash buffer. The beads were further washed for 30 min at RT with 1.5 ml saline buffer (50 mM Tris-HCl, pH 7.5, 150 mM NaCl), containing 0.1% Tween-20 (#9127.1, Roth, Germany), resuspended in 50 µl 2x urea sample buffer (150 mM Tris-HCl, pH 6.8, 6M urea, 6% SDS, 0.01% bromophenol blue) and incubated for 30 min at 1400 rpm and 37°C in a thermomixer. Then, 50 µl boiling buffer (50 mM Tris-HCl, pH 7.5, 1 mM EDTA, 1%, 20% glycerol) was added and the sample was further incubated for 30 min at 1400 rpm and 42°C in a thermomixer. The resulting eluates were subjected to WB analysis. For mass spectrometry analysis of Mup1-GFP 300 OD_600_ equivalents were used with the following modifications: Cells were lysed by with glass beads (3x 1min) at 4°C in 3ml RIPA buffer (50 mM Tris/HCl, pH 7,5, 200 mM NaCl, 10% glycerol, 1% NP-40, 1% Na-Deoxycholate, 0,1% SDS, 2 mM EDTA, 0,1% Tween-20, 25mM NaF, 1x PhosStop (#PHOSS-RO, Sigma, Germany), 1x Complete EDTA-free protease inhibitors, 0.67 mM DTT, 1x yPIC, 2mM PMSF, 20mM NEM. Subsequently another 3 ml of RIPA buffer were added and the lysate was sonicated 5x 1min at 4°C in a water bath sonicator. Lysates were incubated 30 minutes on ice and then centrifuged at 10000g for 10 minutes to remove debris. 80 ul of equilibrated GFP-trap beads were added to the supernatant and incubated rolling for 16 hours at 4°C. Beads were washed at 4°C 3 x 15 min with RIPA buffer supplemented with 500 mM NaCl and then 3x 15 min with urea wash buffer (50 mM Tris/HCl pH 7.5, 100 mM NaCl, 4M Urea). Mup1-GFP was eluated from beads using urea sample buffer as described above. Eluates were separated via SDS-PAGE and Mup1-GFP was visualized by Coomassie staining. Slices were cut from the gel (including the visible Mup1-GFP protein bands and the region above containing ubiquitinated Mup1-GFP) and subjected for further mass spectrometry sample preparation.

### Mass spectrometry sample preparation and analysis

Coomassie-stained gel bands were excised from SDS-PAGE gels, reduced with dithiothreitol, alkylated with iodoacetamide and digested with trypsin (Promega) as previously described (Faserl et al., 2019). Tryptic digest were analyzed using an UltiMate 3000 RSCLnano-HPLC system coupled to a Q Exactive HF mass spectrometer (both Thermo Scientific, Bremen, Germany) equipped with a Nanospray Flex ionization source. The peptides were separated on a homemade fritless fused-silica micro-capillary column (75 µm i.d. x 280 µm o.d. x 10 cm length) packed with 3.0 µm reversed-phase C18 material. Solvents for HPLC were 0.1% formic acid (solvent A) and 0.1% formic acid in 85% acetonitrile (solvent B). The gradient profile was as follows: 0-4 min, 4% B; 4-57 min, 4-35% B; 57-62 min, 35-100% B, and 62-67 min, 100 % B. The flow rate was 250 nL/min.

The Q Exactive HF mass spectrometer was operating in the data dependent mode selecting the top 20 most abundant isotope patterns with charge >1 from the survey scan with an isolation window of 1.6 mass-to-charge ratio (m/z). Survey full scan MS spectra were acquired from 300 to 1750 m/z at a resolution of 60,000 with a maximum injection time (IT) of 120 ms, and automatic gain control (AGC) target 1e6. The selected isotope patterns were fragmented by higher-energy collisional dissociation with normalized collision energy of 28 at a resolution of 30,000 with a maximum IT of 120 ms, and AGC target 5e5.

Data Analysis was performed using Proteome Discoverer 4.1 (Thermo Scientific) with search engine Sequest. The raw files were searched against yeast database (orf_trans_all) with sequence of Mup1-GFP added. Precursor and fragment mass tolerance was set to 10 ppm and 0.02 Da, respectively, and up to two missed cleavages were allowed. Carbamidomethylation of cysteine was set as static modification. Oxidation of methionine, ubiquitination of lysine, and phosphorylation of serine threonine, and tyrosine were set as variable modifications. Peptide identifications were filtered at 1% false discovery rate.

### RNA isolation and quantitative PCR (RT-qPCR)

Exponentially growing or starved cells (40 OD_600nm_) were harvested by centrifugation and immediately frozen in lq. N_2_. Cell pellets were lysed with 1 mm glass beads in a FastPrep-24 homogenizer (MP biosciences) in Qiagen RLT buffer, and RNA was extracted using the RNeasy Mini Kit (#74104, Qiagen, Germany). Yield and purity were determined photometrically. cDNA was prepared from 5 µg DNAse I-treated RNA using the RevertAid First Strand cDNA Synthesis Kit (#K1622, Thermo Fisher, USA) with oligo-dT primer according to the standard protocol. qPCR was performed in 10µl scale with 4µl of cDNA, 5µl TaqMan Gene Expression Master Mix (#4369016, Thermo Fisher, USA) and 0.5 µl TaqMan probe on a PikoReal 96 Real-Time PCR System (Thermo Fisher, USA) with 7 min initial denaturation (95°C) and 40 cycles of 5 sec 95°C and 30 sec 60°C. TaqMan gene expression assays were from Thermo Fisher (*PGK1*: Sc04104844_s1; *ECM21/ART2:* Sc04099967_s1). All probes and primer anneal within coding sequences. Each RT-qPCR analysis was done from 2-3 independent biological samples in 3-4 technical replicates. Data were analyzed with the PikoReal software (version 2.2; Thermo Fisher) with manual threshold adjustment, and relative mRNA abundance was calculated in Microsoft Excel using the ΔΔC_T_ method. Statistical comparisons were calculated using the Student t-test.

## Acknowledgments

We are greatful to Claudine Kraft and Scott Emr for reagents. This work was supported by EMBO/Marie Curie (ALTF 642-2012; EMBOCOFUND2010, GA-2010-267146) and ‘Tiroler Wissenschaftsfond’ to O.S., Austrian Science Fund (FWF-Y444-B12, P30263, P29583) and MCBO (W1101-B18) to D.T and Agence Nationale pour la Recherche (“P-Nut”, ANR-16-CE13-0002-01) to S.L.

## Author contributions

V.I., J.Z., S.W., J.K, R.G., T.J., G.K.B and L.A.H. performed the screens. V.I. and J.Z. analyzed Mup1 ubiquitination and J.Z. characterized the contribution of the GAAC. V.I and S.S. analyzed Mup1 degrons and Art2 basic patches. V.I., L.K. and H.L performed and analyzed mass spectrometry experiments. V.I., J.Z., S.S., J.K., S.L. and O.S. constructed yeast strains and plasmids and conducted microscopy, biochemical and genetic experiments. O.S. and D.T. and analyzed data, conceptualized and guided the study; S.L., D.T. and O.S. wrote the manuscript.

---

**Figure S1:**
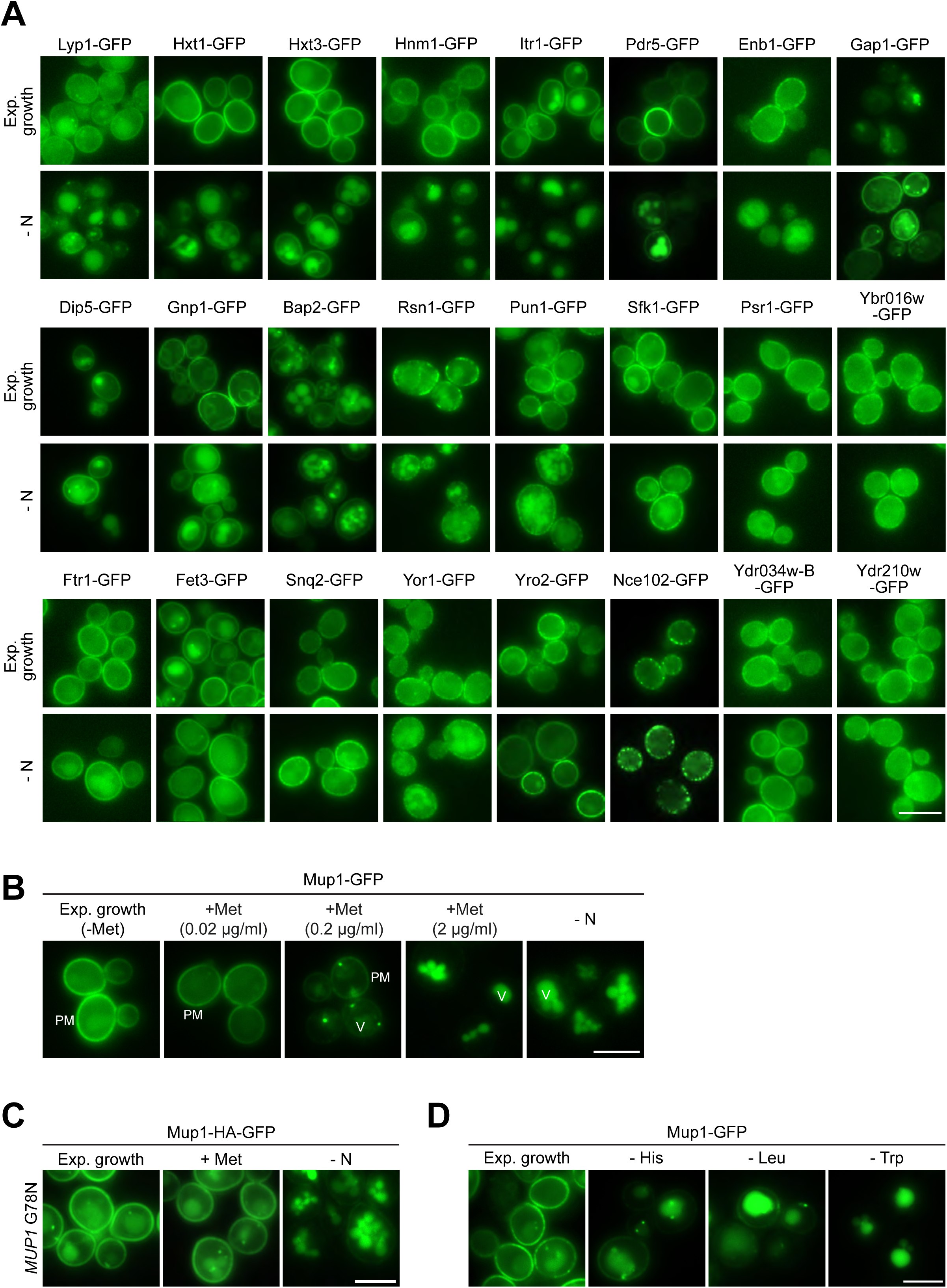
**Related to Fig. 1 A)** Live-cell fluorescence microscopy analysis of chromosomally GFP-tagged plasma membrane proteins. Cells were starved (- N) for 6-8h after 24h exponential growth. **B)** Live-cell fluorescence microscopy analysis of WT cells expressing *MUP1-GFP*. Cells were treated with the indicated methionine concentrations (+ Met) for 1.5h or starved (- N) for 6h after 24h exponential growth. **C)** Live-cell fluorescence microscopy analysis of *MUP1(G78N)-HA-GFP* cells. Cells were treated with 20 µg/ml L-methionine (+ Met) for 1.5h or starved (- N) for 6h after 24h exponential growth. PM: plasma membrane; V: vacuole. Scale bars = 5 µm. **D)** Live-cell fluorescence microscopy analysis of WT cells expressing *MUP1-GFP*. Cells were starved for histidine (- His), leucine (- Leu) or tryptophan (- Trp) for 4.5h after in presence of all other amino acids required for growth after 24h nutrient replete exponential growth.

**Figure. S2:**
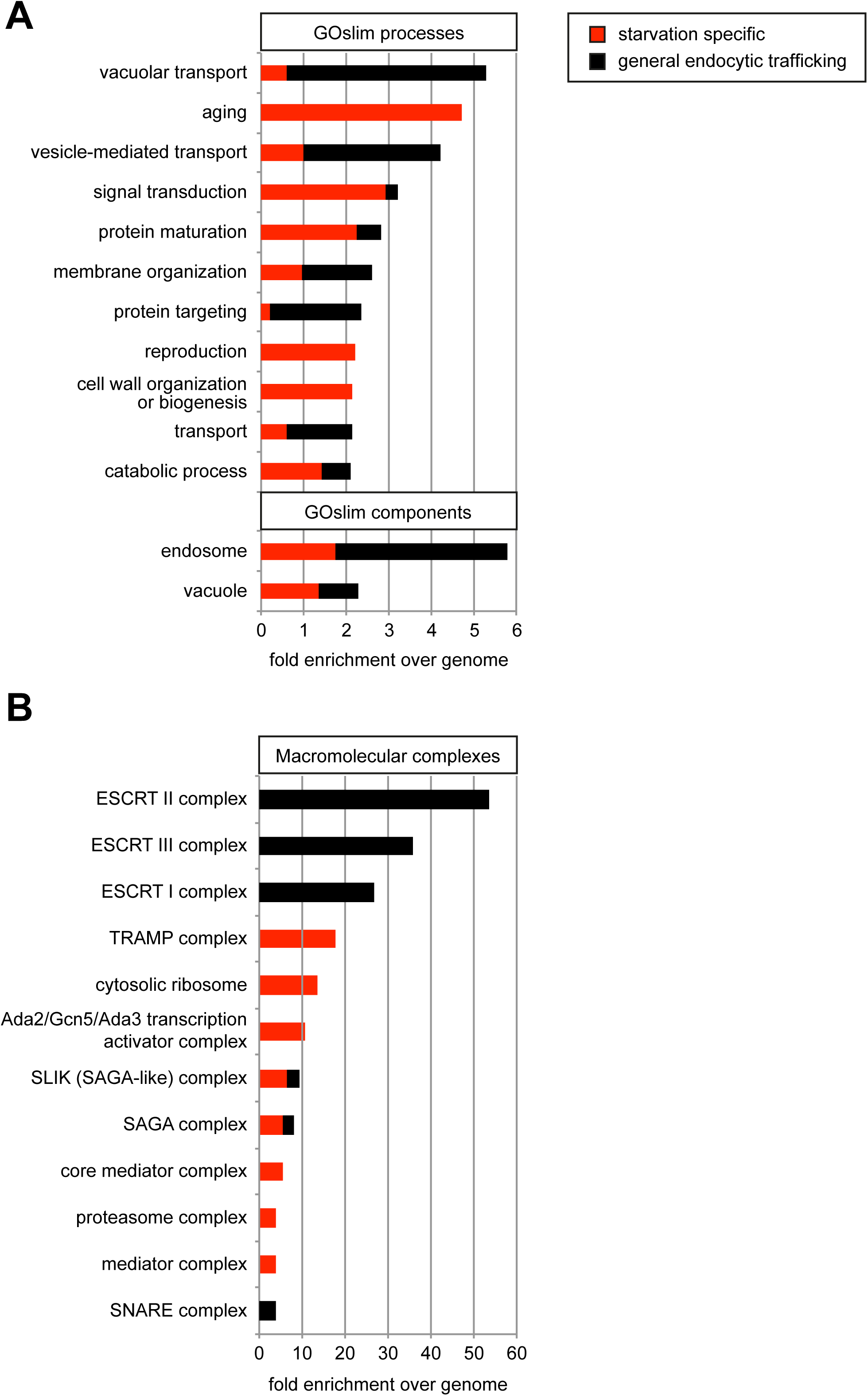
**related to Fig. 2 A)** Enrichment of cellular processes and components GO terms in 128 hits from the genome-wide screen for genes involved in the starvation-induced endocytosis of Mup1-pHluorin (table S2). Data are represented as fold-enrichment over whole genome frequency, with the fraction of starvation-specific genes in red and general endocytic trafficking regulators in black. Only GO terms with more than two-fold enrichment over genome were included. (B) Enrichment of macromolecular complex GO terms in 128 hits from the genome-wide screen for genes involved in the starvation-induced endocytosis of Mup1-pHluorin analyzed as in (A). See also table S3.

**Figure S3:**
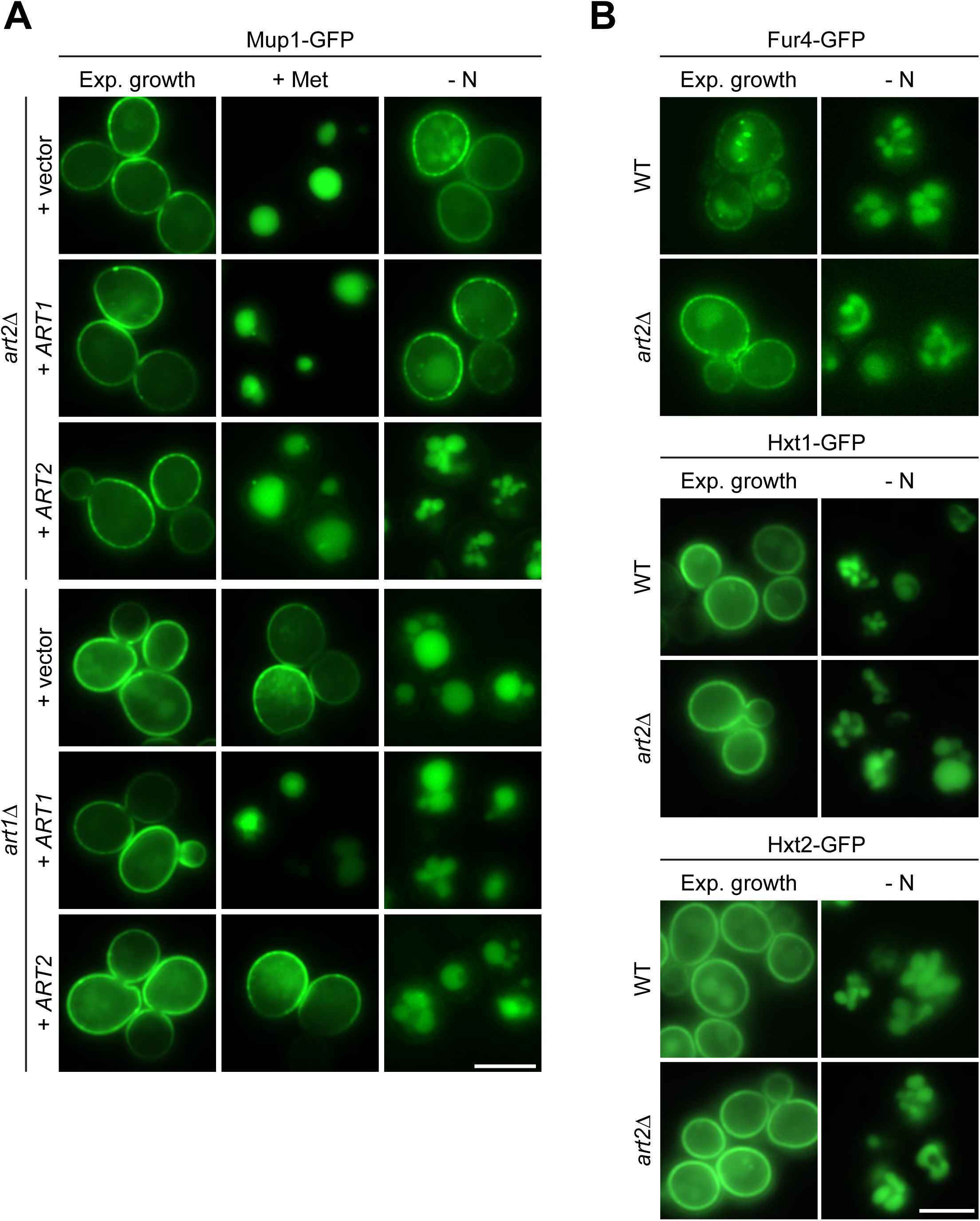
**Related to Fig. 3 A)** Live-cell fluorescence microscopy analysis of *art2*Δ and *art1Δ* cells expressing *MUP1-GFP* and plasmids encoding *ART1*, *ART2* or empty vectors as indicated. Cells were treated with 20 µg/ml L-methionine (+ Met) for 1.5h or starved (- N) for 6h after 24h exponential growth. **B)** Live-cell fluorescence microscopy analysis of wild type (WT) and *art2*Δ cells expressing pRS416*-FUR4-GFP*, *HXT1-GFP* or *HXT2-GFP*. Cells were starved (- N) for 6h after 24h exponential growth. Scale bars = 5 µm.

**Figure S4:**
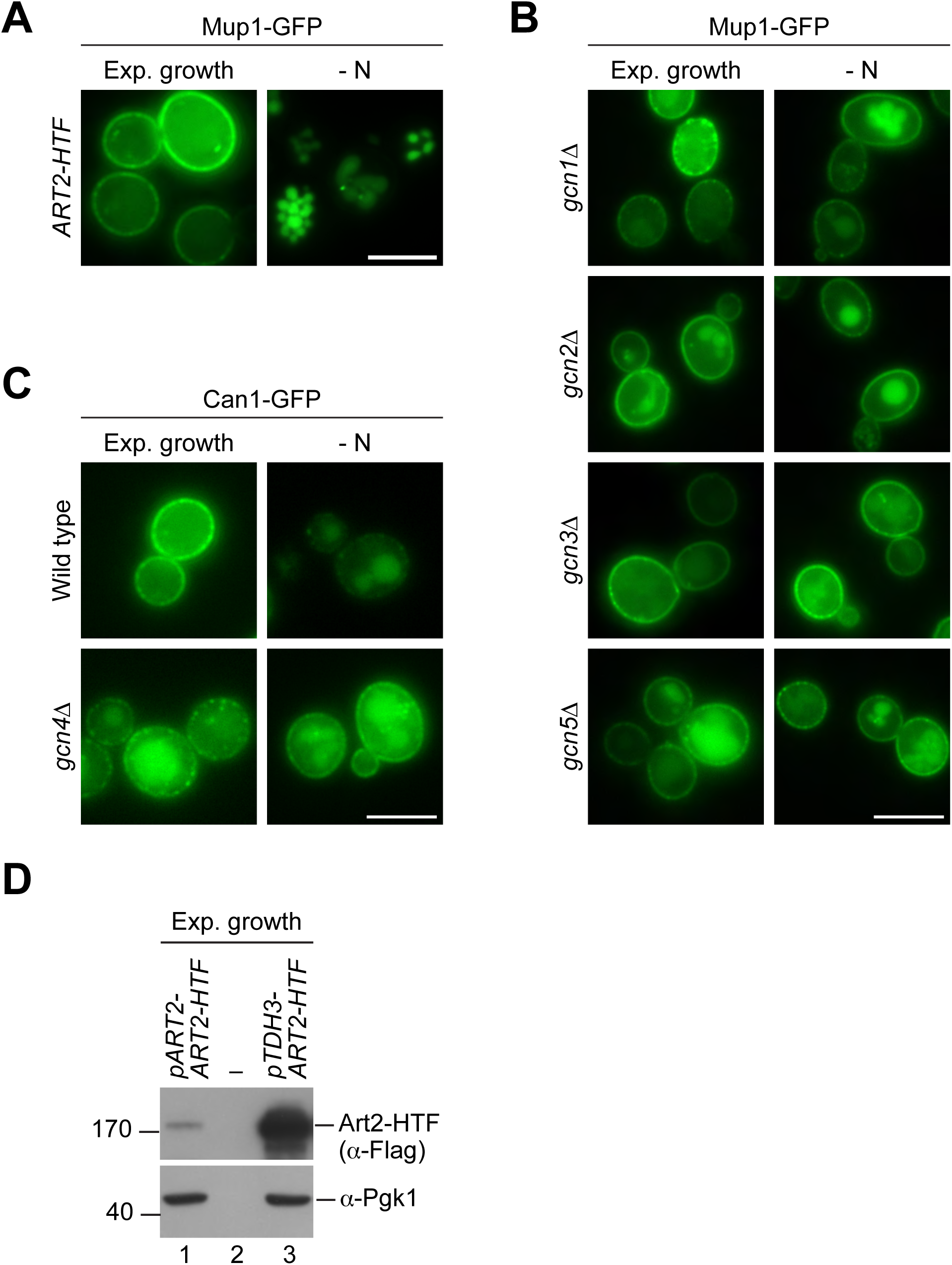
**Related to Fig. 4. A)-C)** Live-cell fluorescence microscopy analysis of the indicated strains starved (- N) for 6h after 24h exponential growth. **D)** SDS PAGE and Western blot analysis with the indicated antibodies of whole cell protein extracts from WT cells expressing *pART2-ART2-HTF* or pRS416-*pTDH3-ART2-HTF* after 24h exponential growth. Scale bars = 5 µm.

**Figure S5:**
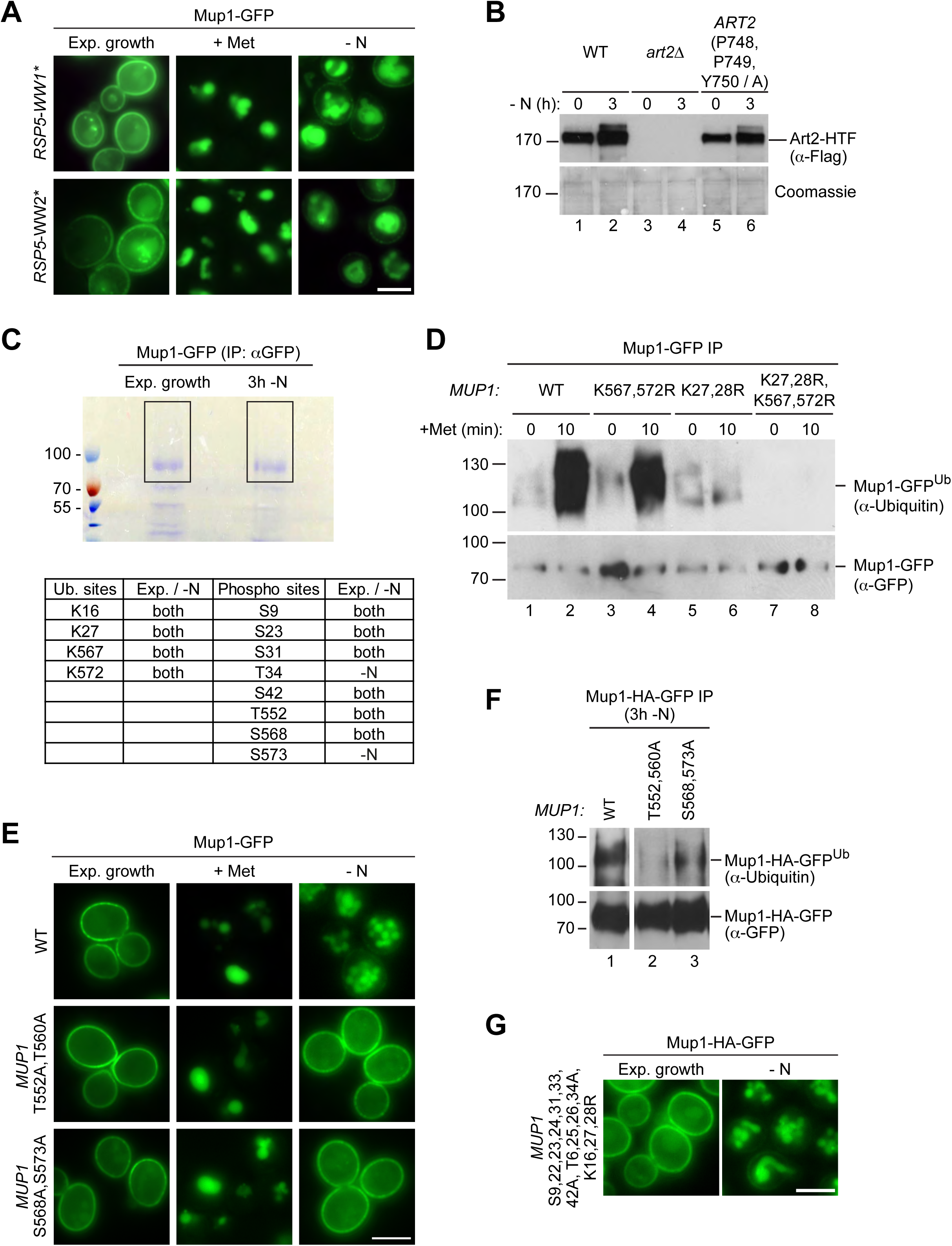
**Related to Fig. 5 A)** Live-cell fluorescence microscopy analysis of *rsp5*Δ cells expressing pRS416-*MUP1-GFP* and pRS415-*HTF-RSP5-WW1\** or pRS415-*HTF-RSP5-WW2\**. Cells were treated with 20 µg/ml L-methionine (+ Met) for 1.5h or starved (- N) for 6h after 24h exponential growth. **B)** SDS PAGE and Western blot analysis with the indicated antibodies of whole cell protein extracts from *art2Δ* cells expressing pRS416-*ART2-HTF* (WT), pRS416 or pRS416-*ART2 P748A,P749A,Y750A-HTF*. Cells were starved (- N) for 3h after 24h exponential growth. **C)** Upper panel: Coomassie-stained SDS PAGE of immunoprecipitated Mup1-GFP from WT cells after 24h exponential growth or subsequent 3h starvation (-N). The black rectangles indicate the regions of the gel analyzed by mass spectrometry. Lower panel: Ubiquitinated lysine (K) and phosphorylated serine (S) and threonine (T) residues of Mup1 identified by mass spectrometry during exponential growth and/or starvation with numbers corresponding to amino acid positions in the Mup1 sequence. **D)** SDS PAGE and Western blot analysis with the indicated antibodies of immunoprecipitated Mup1-GFP from WT cells or the indicated *MUP1* mutants treated with 20 µg/ml L-methionine for 10 min after 24h of exponential growth. Equal amounts of immunoprecipitated Mup1-GFP were loaded to compare the extent of ubiquitination. **E)** Live-cell fluorescence microscopy analysis of cells expressing *MUP1-GFP* (wild type (WT)), *MUP1 T552,560A-GFP* or *MUP1 S568,573A-GFP*. Cells were treated with 20 µg/ml L-methionine (+ Met) for 1.5h or starved (- N) for 6h after 24h exponential growth. **F)** *MUP1-HA-GFP* (wild type (WT)), *MUP1 T552,560A-HA-GFP* or *MUP1 S568,573A-HA-GFP* cells were starved (- N) for 3h and analyzed as in D). **G)** Live-cell fluorescence microscopy analysis of cells expressing *MUP1 T6,25,26,34A,S9,22,23,24,31,33,42A,K16,27,28R-HA-GFP*. Cells were starved (- N) for 3h after 24h exponential growth. Scale bars = 5 µm.

**Figure S6:**
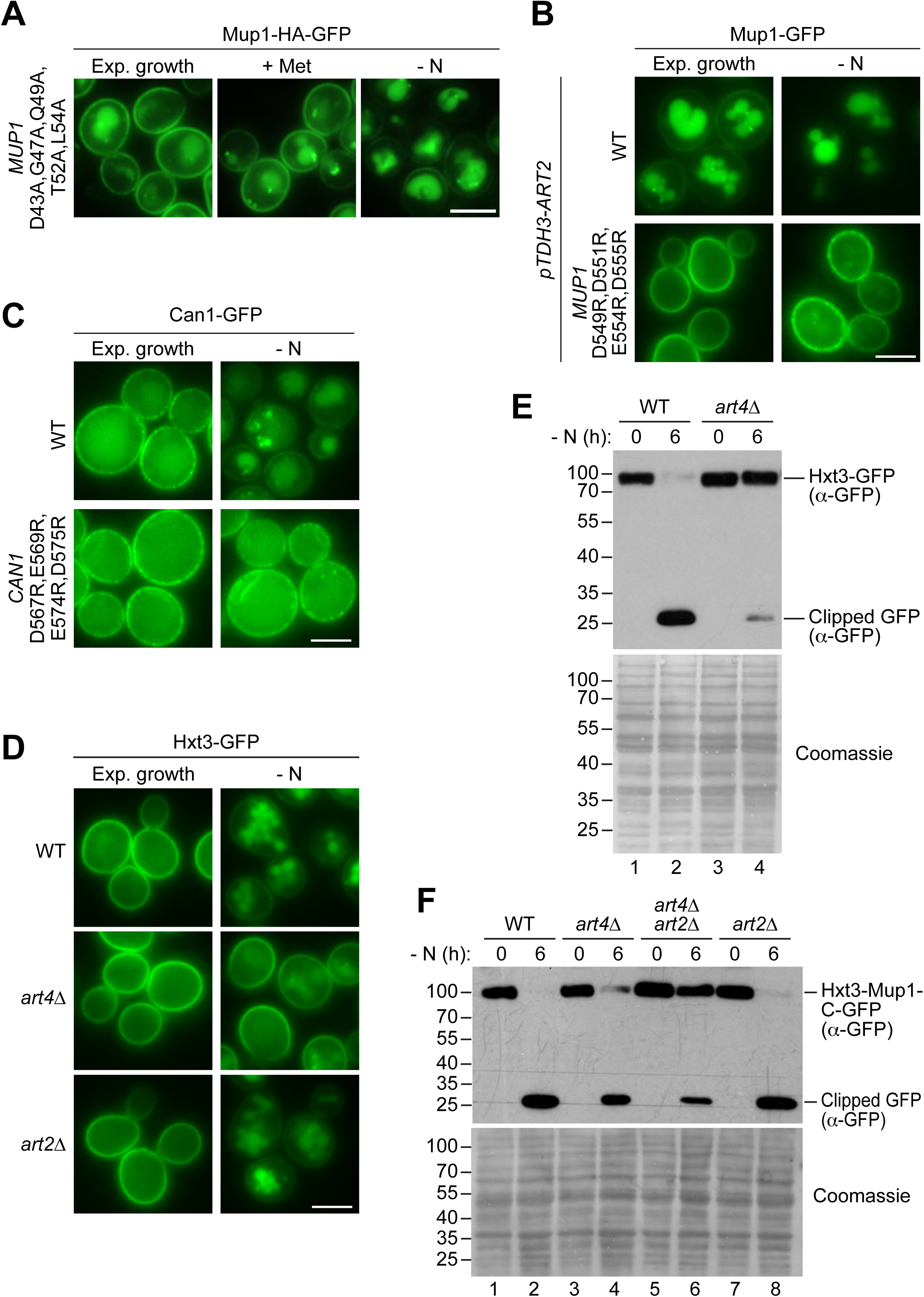
**Related to Fig. 6 A)-D)** Live-cell fluorescence microscopy analysis of the indicated strains treated with 20 µg/ml L-methionine (+ Met) for 1.5h or starved (- N) for 6h after 24h exponential growth. **E)** SDS PAGE and Western blot analysis with the indicated antibodies of whole cell protein extracts from wild type (WT) and *art4*Δ cells expressing *HXT3-GFP*. Cells were starved (- N) for 6h after 24h exponential growth. Coomassie staining of the membrane serves as loading control. **F)** Wild type (WT), *art4*Δ, *art2*Δ *art4*Δ and *art2*Δ cells expressing *HXT3-MUP1-C-GFP* were treated as in E). Scale bars = 5 µm.

**Figure S7:**
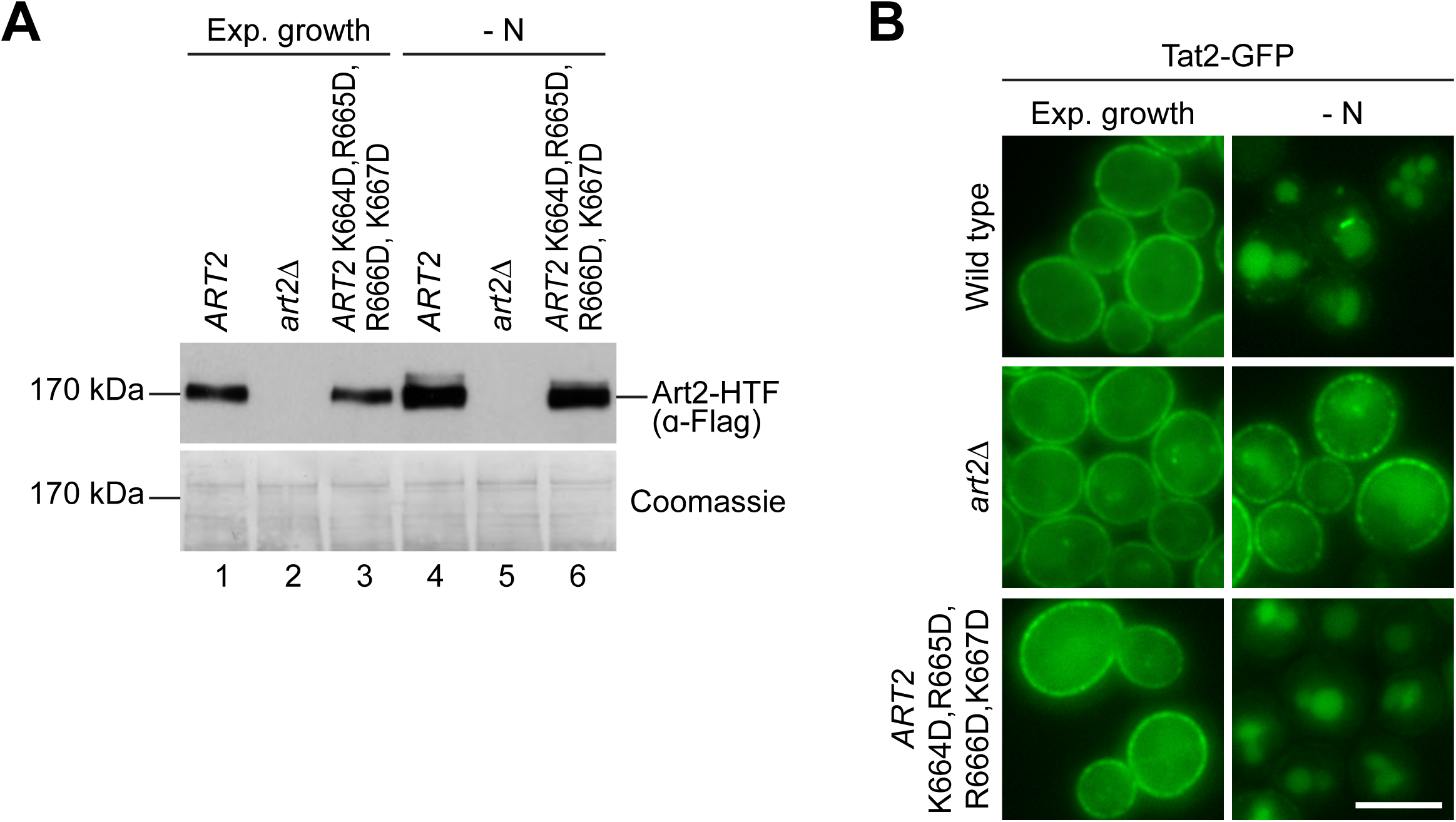
**Related to Fig. 7 A)** SDS PAGE and Western blot analysis with the indicated antibodies of whole cell protein extracts from *art2*Δ cells expressing pRS416-*ART2-HTF*, empty vector or pRS416-*ART2 K664D,R665D,R666D,K667D-HTF*. Cells were starved (- N) for 3h after 24h exponential growth. **B)** Live-cell fluorescence microscopy analysis of *art2*Δ cells expressing *TAT2-GFP* and *pRS416-ART2*, empty vector or *pRS416-ART2 K664D,R665D,R666D,K667D*. Cells were starved (- N) for 6h after 24h exponential growth. Scale bars = 5 µm.

**Table S1:**
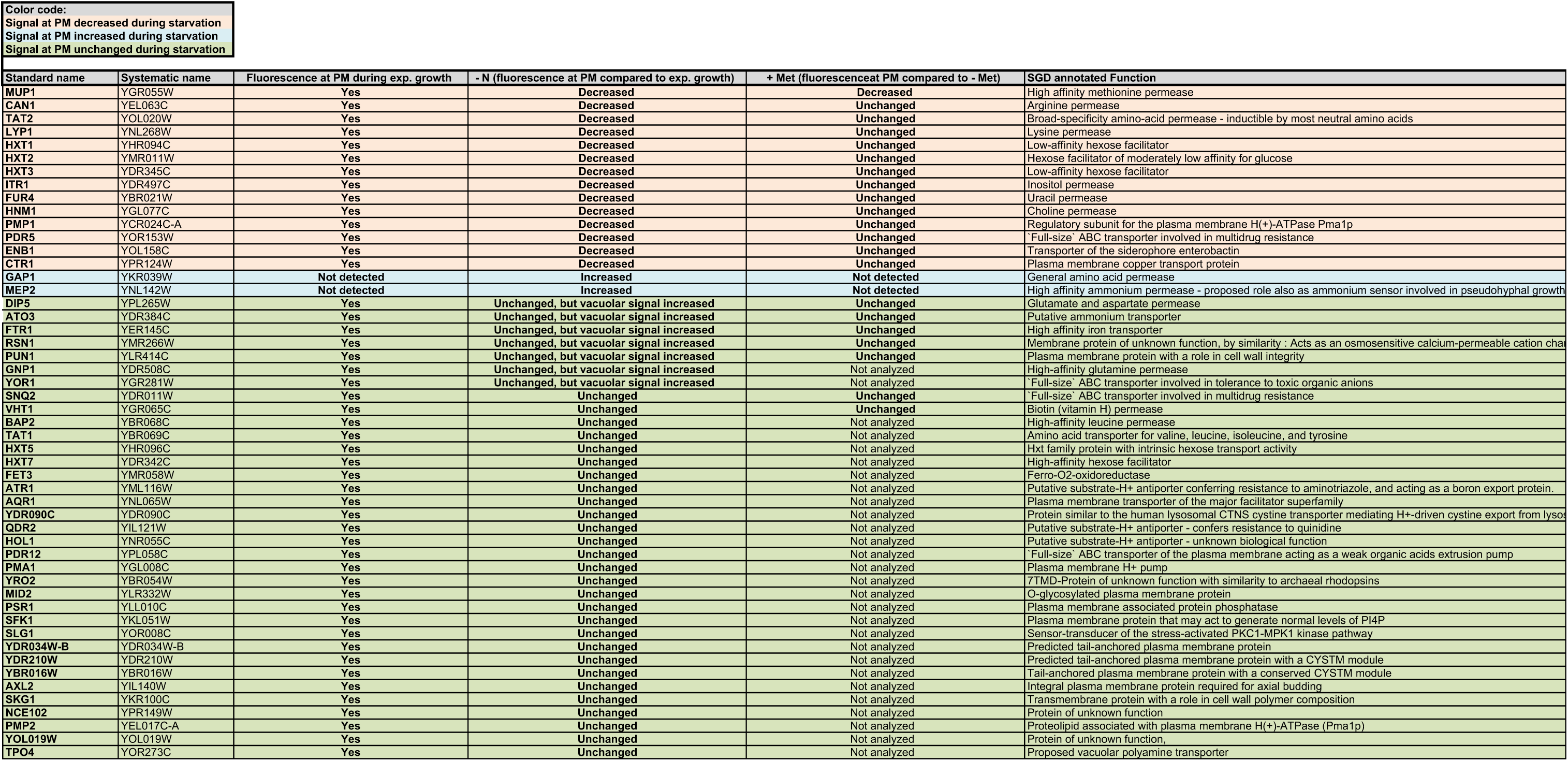
Fluorescence microscopy-based screen for substrates of starvation-induced endocytosis.

**Table S2:**
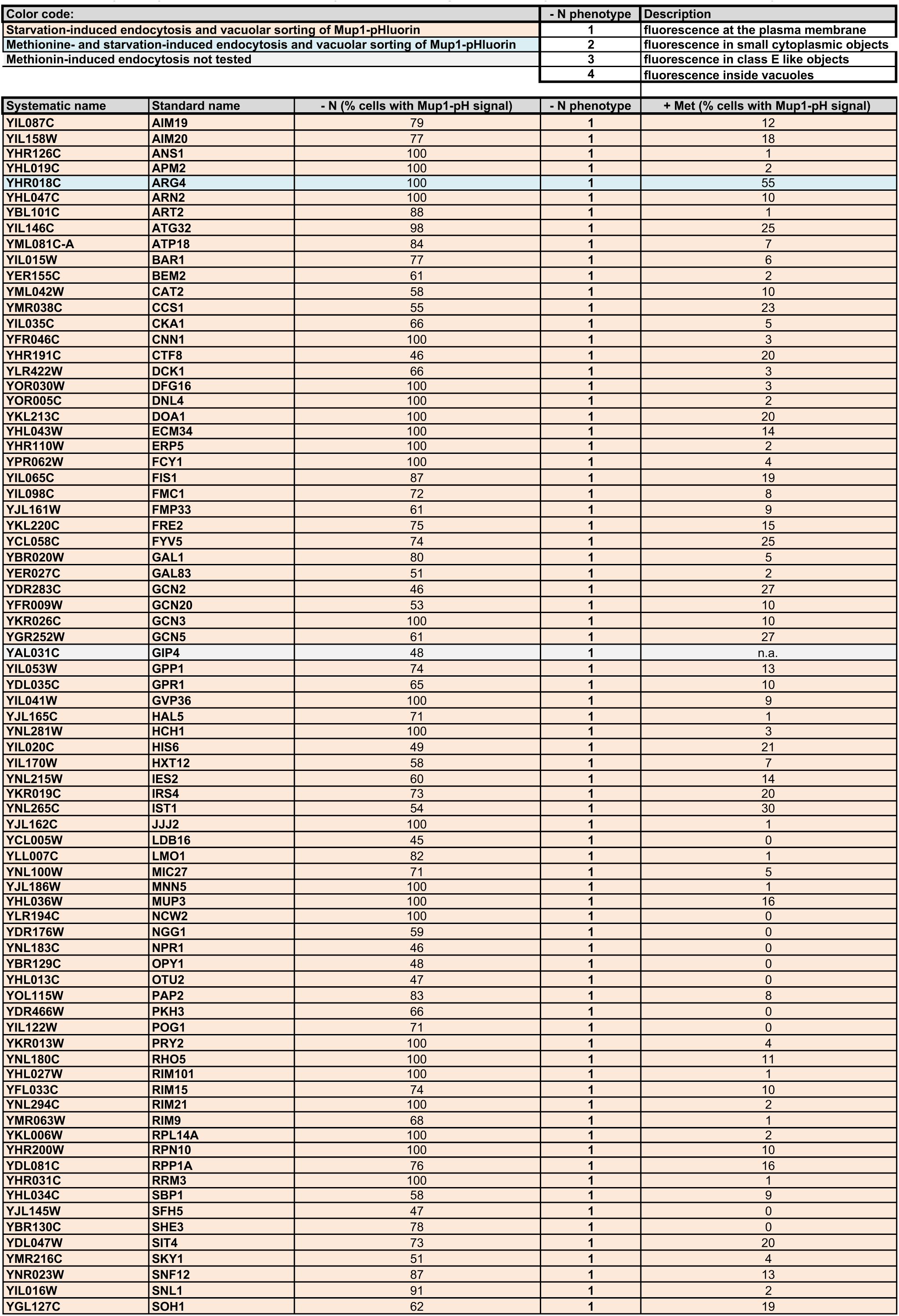
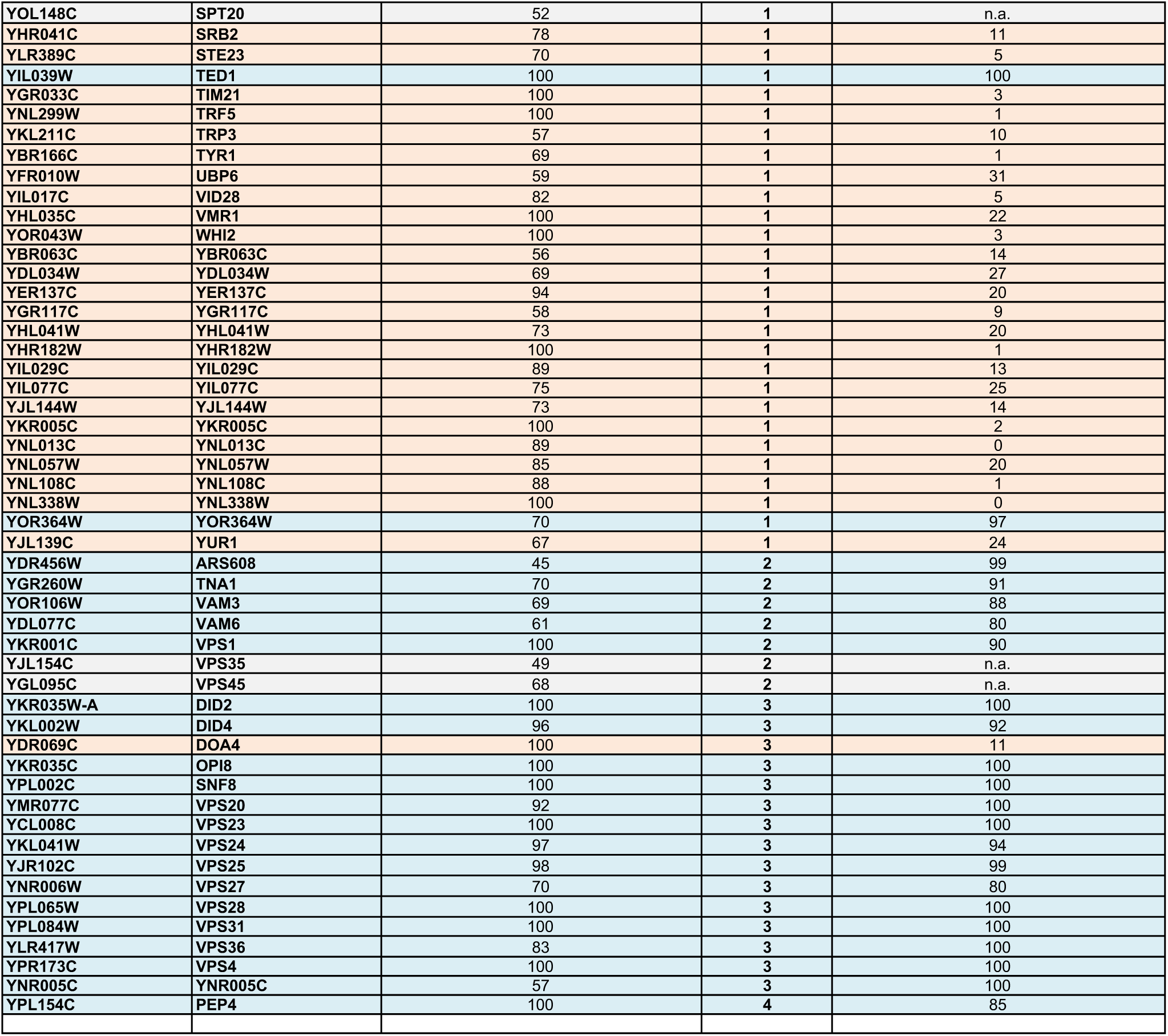
Flow cytometry and fluorescence microscopy-based screen for genes specifically involved in starvation-induced endocytosis Description.

| Color code: |  | - N phenotype | Description |
| --- | --- | --- | --- |
| Starvation-induced endocytosis and vacuolar sorting of Mup1-pHluorin |  | 1 | fluorescence at the plasma membrane |
| Methionine- and starvation-induced endocytosis and vacuolar sorting of Mup1-pHluorin |  | 2 | fluorescence in small cytoplasmic objects |
| Methionin-induced endocytosis not tested |  | 3 | fluorescence in class E like objects |
|  |  | 4 | fluorescence inside vacuoles |

| Systematic name | Standard name | - N (% cells with Mup1-pH signal) | - N phenotype | + Met (% cells with Mup1-pH signal) |
| --- | --- | --- | --- | --- |
| YIL087C | AIM19 | 79 | 1 | 12 |
| YIL158W | AIM20 | 77 | 1 | 18 |
| YHR126C | ANS1 | 100 | 1 | 1 |
| YHL019C | APM2 | 100 | 1 | 2 |
| YHR018C | ARG4 | 100 | 1 | 55 |
| YHL047C | ARN2 | 100 | 1 | 10 |
| YBL101C | ART2 | 88 | 1 | 1 |
| YIL146C | ATG32 | 98 | 1 | 25 |
| YML081C-A | ATP18 | 84 | 1 | 7 |
| YIL015W | BAR1 | 77 | 1 | 6 |
| YER155C | BEM2 | 61 | 1 | 2 |
| YML042W | CAT2 | 58 | 1 | 10 |
| YMR038C | CCS1 | 55 | 1 | 23 |
| YIL035C | CKA1 | 66 | 1 | 5 |
| YFR046C | CNN1 | 100 | 1 | 3 |
| YHR191C | CTF8 | 46 | 1 | 20 |
| YLR422W | DCK1 | 66 | 1 | 3 |
| YOR030W | DFG16 | 100 | 1 | 3 |
| YOR005C | DNL4 | 100 | 1 | 2 |
| YKL213C | DOA1 | 100 | 1 | 20 |
| YHL043W | ECM34 | 100 | 1 | 14 |
| YHR110W | ERP5 | 100 | 1 | 2 |
| YPR062W | FCY1 | 100 | 1 | 4 |
| YIL065C | FIS1 | 87 | 1 | 19 |
| YIL098C | FMC1 | 72 | 1 | 8 |
| YJL161W | FMP33 | 61 | 1 | 9 |
| YKL220C | FRE2 | 75 | 1 | 15 |
| YCL058C | FYV5 | 74 | 1 | 25 |
| YBR020W | GAL1 | 80 | 1 | 5 |
| YER027C | GAL83 | 51 | 1 | 2 |
| YDR283C | GCN2 | 46 | 1 | 27 |
| YFR009W | GCN20 | 53 | 1 | 10 |
| YKR026C | GCN3 | 100 | 1 | 10 |
| YGR252W | GCN5 | 61 | 1 | 27 |
| YAL031C | GIP4 | 48 | 1 | n.a. |
| YIL053W | GPP1 | 74 | 1 | 13 |
| YDL035C | GPR1 | 65 | 1 | 10 |
| YIL041W | GVP36 | 100 | 1 | 9 |
| YJL165C | HAL5 | 71 | 1 | 1 |
| YNL281W | HCH1 | 100 | 1 | 3 |
| YIL020C | HIS6 | 49 | 1 | 21 |
| YIL170W | HXT12 | 58 | 1 | 7 |
| YNL215W | IES2 | 60 | 1 | 14 |
| YKR019C | IRS4 | 73 | 1 | 20 |
| YNL265C | IST1 | 54 | 1 | 30 |
| YJL162C | JJJ2 | 100 | 1 | 1 |
| YCL005W | LDB16 | 45 | 1 | 0 |
| YLL007C | LMO1 | 82 | 1 | 1 |
| YNL100W | MIC27 | 71 | 1 | 5 |
| YJL186W | MNN5 | 100 | 1 | 1 |
| YHL036W | MUP3 | 100 | 1 | 16 |
| YLR194C | NCW2 | 100 | 1 | 0 |
| YDR176W | NGG1 | 59 | 1 | 0 |
| YNL183C | NPR1 | 46 | 1 | 0 |
| YBR129C | OPY1 | 48 | 1 | 0 |
| YHL013C | OTU2 | 47 | 1 | 0 |
| YOL115W | PAP2 | 83 | 1 | 8 |
| YDR466W | PKH3 | 66 | 1 | 0 |
| YIL122W | POG1 | 71 | 1 | 0 |
| YKR013W | PRY2 | 100 | 1 | 4 |
| YNL180C | RHO5 | 100 | 1 | 11 |
| YHL027W | RIM101 | 100 | 1 | 1 |
| YFL033C | RIM15 | 74 | 1 | 10 |
| YNL294C | RIM21 | 100 | 1 | 2 |
| YMR063W | RIM9 | 68 | 1 | 1 |
| YKL006W | RPL14A | 100 | 1 | 2 |
| YHR200W | RPN10 | 100 | 1 | 10 |
| YDL081C | RPP1A | 76 | 1 | 16 |
| YHR031C | RRM3 | 100 | 1 | 1 |
| YHL034C | SBP1 | 58 | 1 | 9 |
| YJL145W | SFH5 | 47 | 1 | 0 |
| YBR130C | SHE3 | 78 | 1 | 0 |
| YDL047W | SIT4 | 73 | 1 | 20 |
| YMR216C | SKY1 | 51 | 1 | 4 |
| YNR023W | SNF12 | 87 | 1 | 13 |
| YIL016W | SNL1 | 91 | 1 | 2 |
| YGL127C | SOH1 | 62 | 1 | 19 |
| YOL148C | SPT20 | 52 | 1 | n.a. |
| YHR041C | SRB2 | 78 | 1 | 11 |
| YLR389C | STE23 | 70 | 1 | 5 |
| YIL039W | TED1 | 100 | 1 | 100 |
| YGR033C | TIM21 | 100 | 1 | 3 |
| YNL299W | TRF5 | 100 | 1 | 1 |
| YKL211C | TRP3 | 57 | 1 | 10 |
| YBR166C | TYR1 | 69 | 1 | 1 |
| YFR010W | UBP6 | 59 | 1 | 31 |
| YIL017C | VID28 | 82 | 1 | 5 |
| YHL035C | VMR1 | 100 | 1 | 22 |
| YOR043W | WHI2 | 100 | 1 | 3 |
| YBR063C | YBR063C | 56 | 1 | 14 |
| YDL034W | YDL034W | 69 | 1 | 27 |
| YER137C | YER137C | 94 | 1 | 20 |
| YGR117C | YGR117C | 58 | 1 | 9 |
| YHL041W | YHL041W | 73 | 1 | 20 |
| YHR182W | YHR182W | 100 | 1 | 1 |
| YIL029C | YIL029C | 89 | 1 | 13 |
| YIL077C | YIL077C | 75 | 1 | 25 |
| YJL144W | YJL144W | 73 | 1 | 14 |
| YKR005C | YKR005C | 100 | 1 | 2 |
| YNL013C | YNL013C | 89 | 1 | 0 |
| YNL057W | YNL057W | 85 | 1 | 20 |
| YNL108C | YNL108C | 88 | 1 | 1 |
| YNL338W | YNL338W | 100 | 1 | 0 |
| YOR364W | YOR364W | 70 | 1 | 97 |
| YJL139C | YUR1 | 67 | 1 | 24 |
| YDR456W | ARS608 | 45 | 2 | 99 |
| YGR260W | TNA1 | 70 | 2 | 91 |
| YOR106W | VAM3 | 69 | 2 | 88 |
| YDL077C | VAM6 | 61 | 2 | 80 |
| YKR001C | VPS1 | 100 | 2 | 90 |
| YJL154C | VPS35 | 49 | 2 | n.a. |
| YGL095C | VPS45 | 68 | 2 | n.a. |
| YKR035W-A | DID2 | 100 | 3 | 100 |
| YKL002W | DID4 | 96 | 3 | 92 |
| YDR069C | DOA4 | 100 | 3 | 11 |
| YKR035C | OPI8 | 100 | 3 | 100 |
| YPL002C | SNF8 | 100 | 3 | 100 |
| YMR077C | VPS20 | 92 | 3 | 100 |
| YCL008C | VPS23 | 100 | 3 | 100 |
| YKL041W | VPS24 | 97 | 3 | 94 |
| YJR102C | VPS25 | 98 | 3 | 99 |
| YNR006W | VPS27 | 70 | 3 | 80 |
| YPL065W | VPS28 | 100 | 3 | 100 |
| YPL084W | VPS31 | 100 | 3 | 100 |
| YLR417W | VPS36 | 83 | 3 | 100 |
| YPR173C | VPS4 | 100 | 3 | 100 |
| YNR005C | YNR005C | 57 | 3 | 100 |
| YPL154C | PEP4 | 100 | 4 | 85 |

**Table S3:**
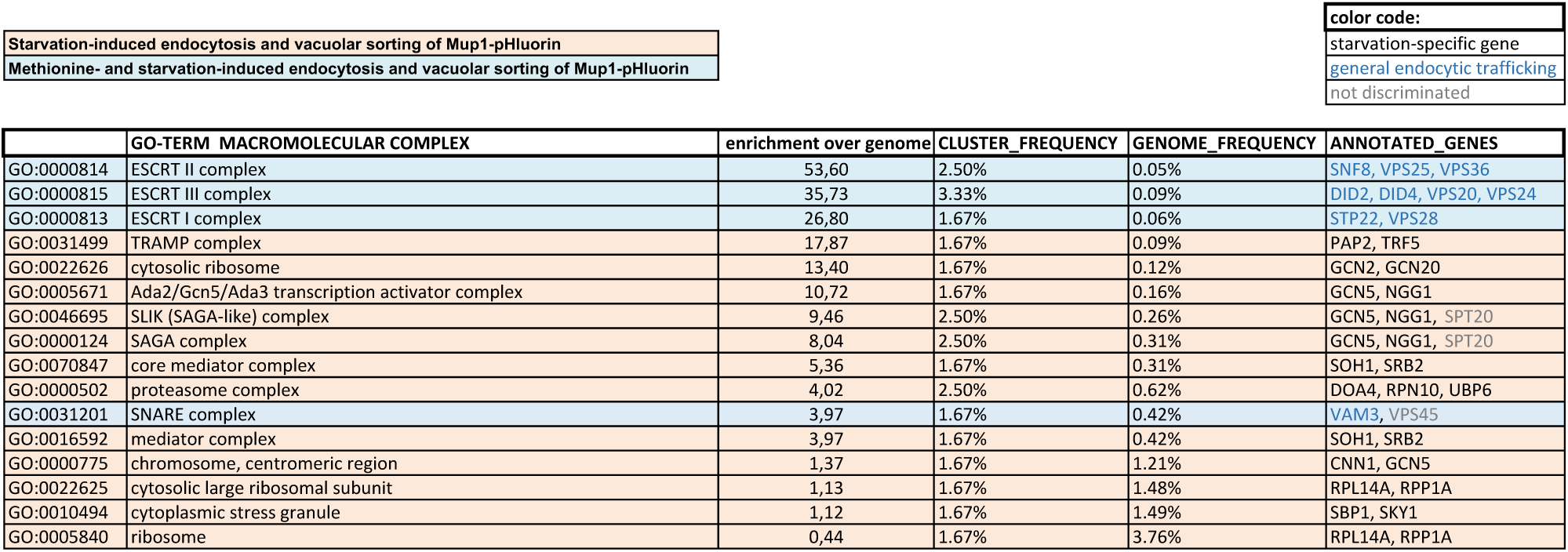
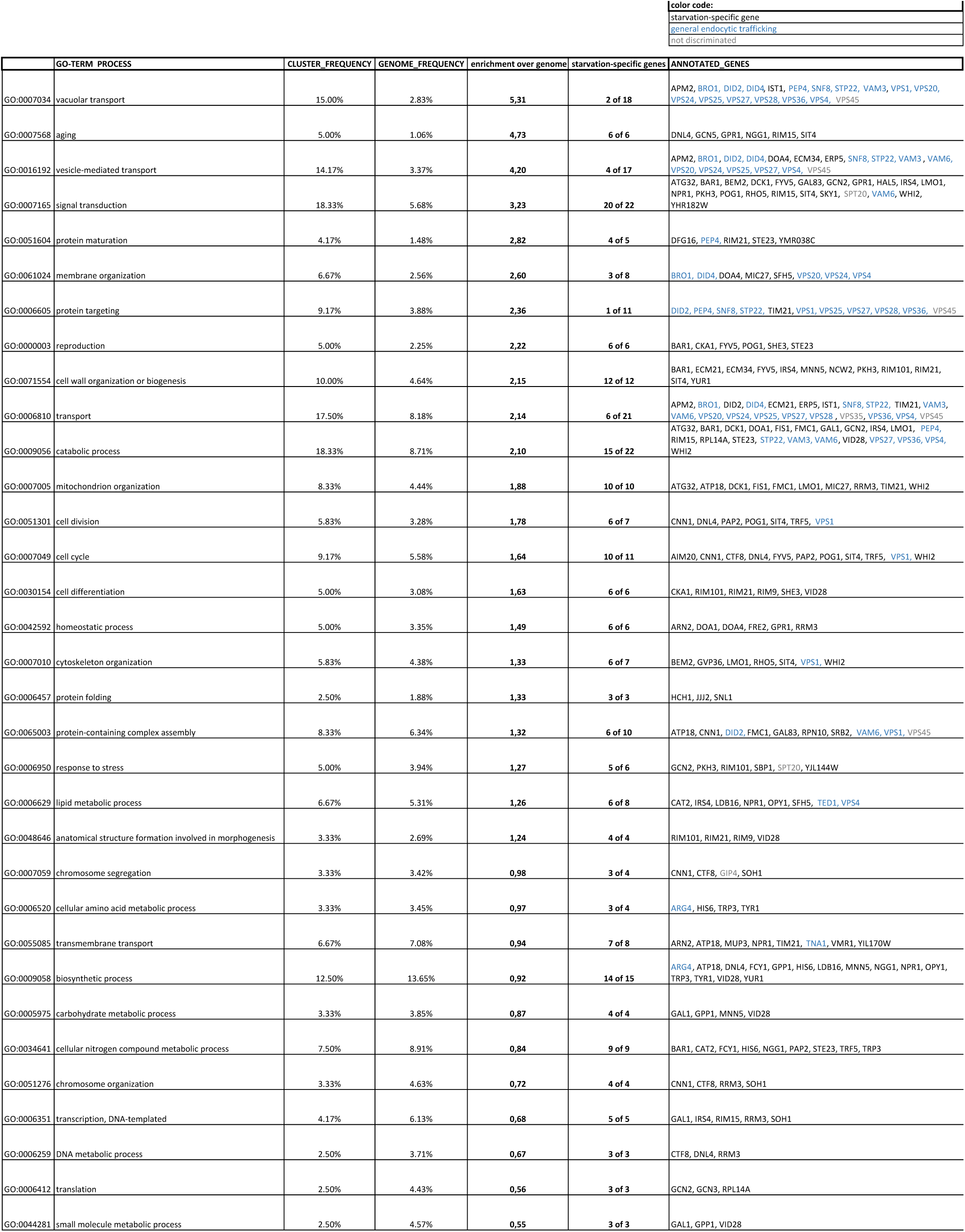
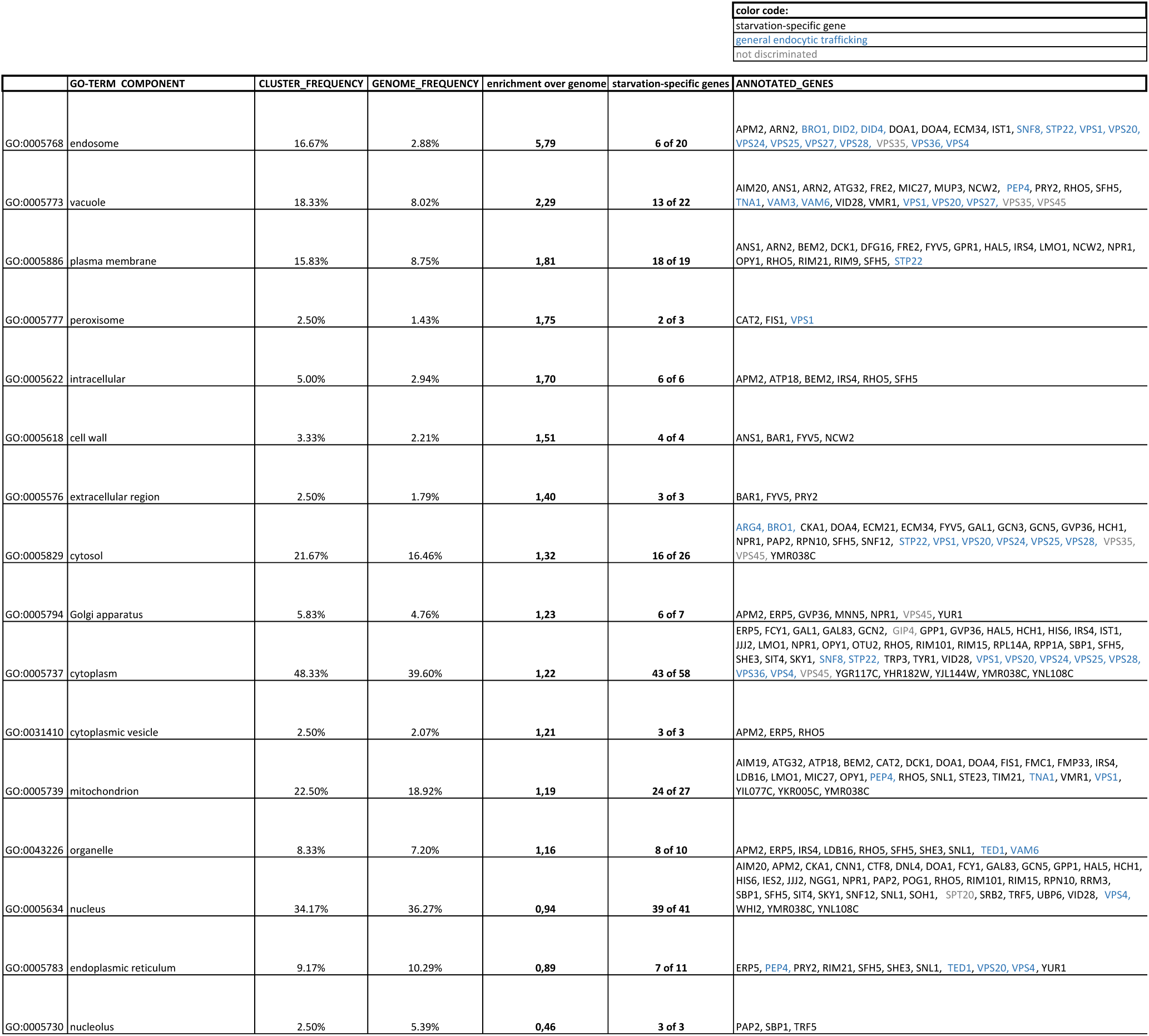
Gene ontology term analysis of 128 genes required for starvation-induced endocytosis of Mup1-pHluorin.

**Table S4:**
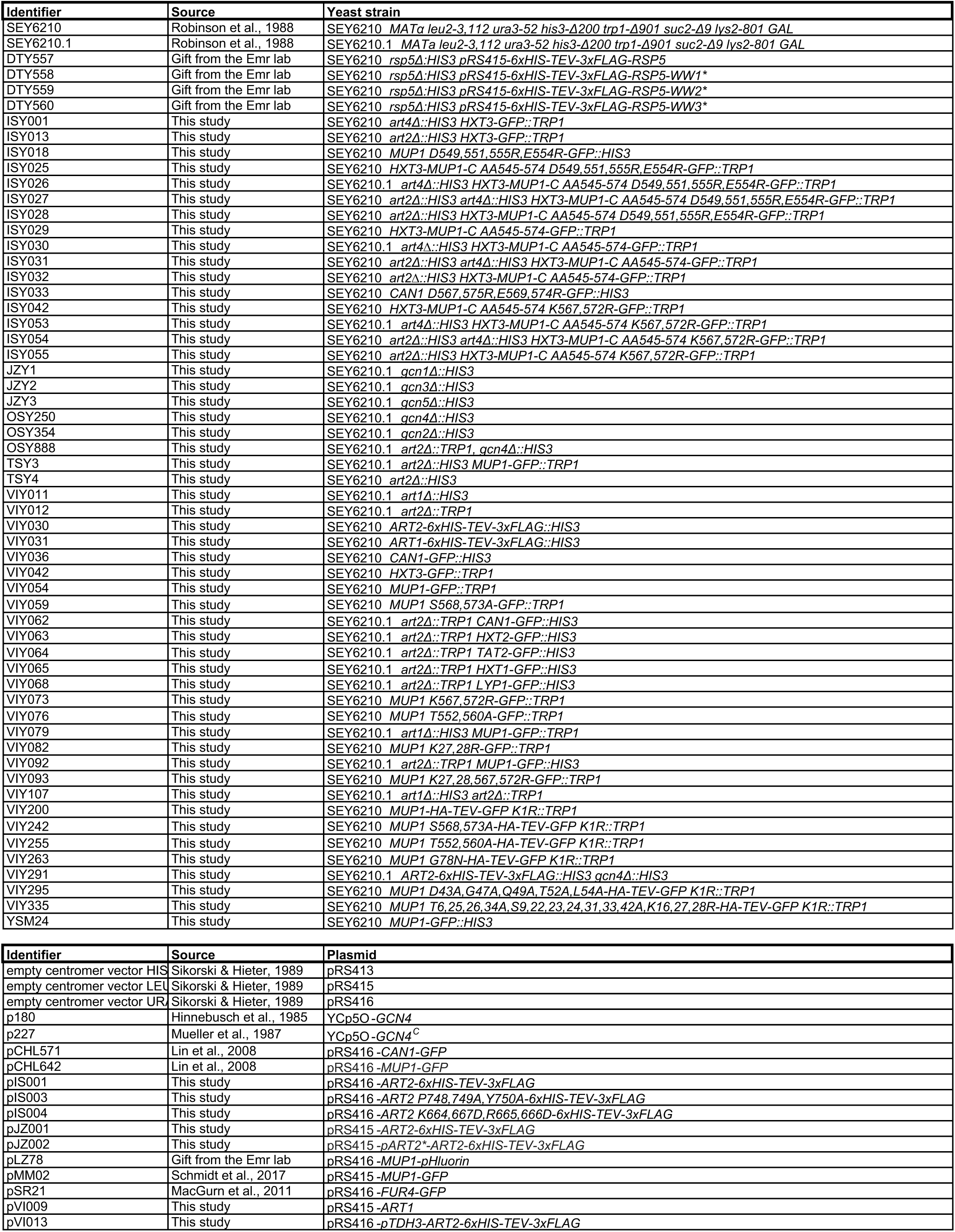
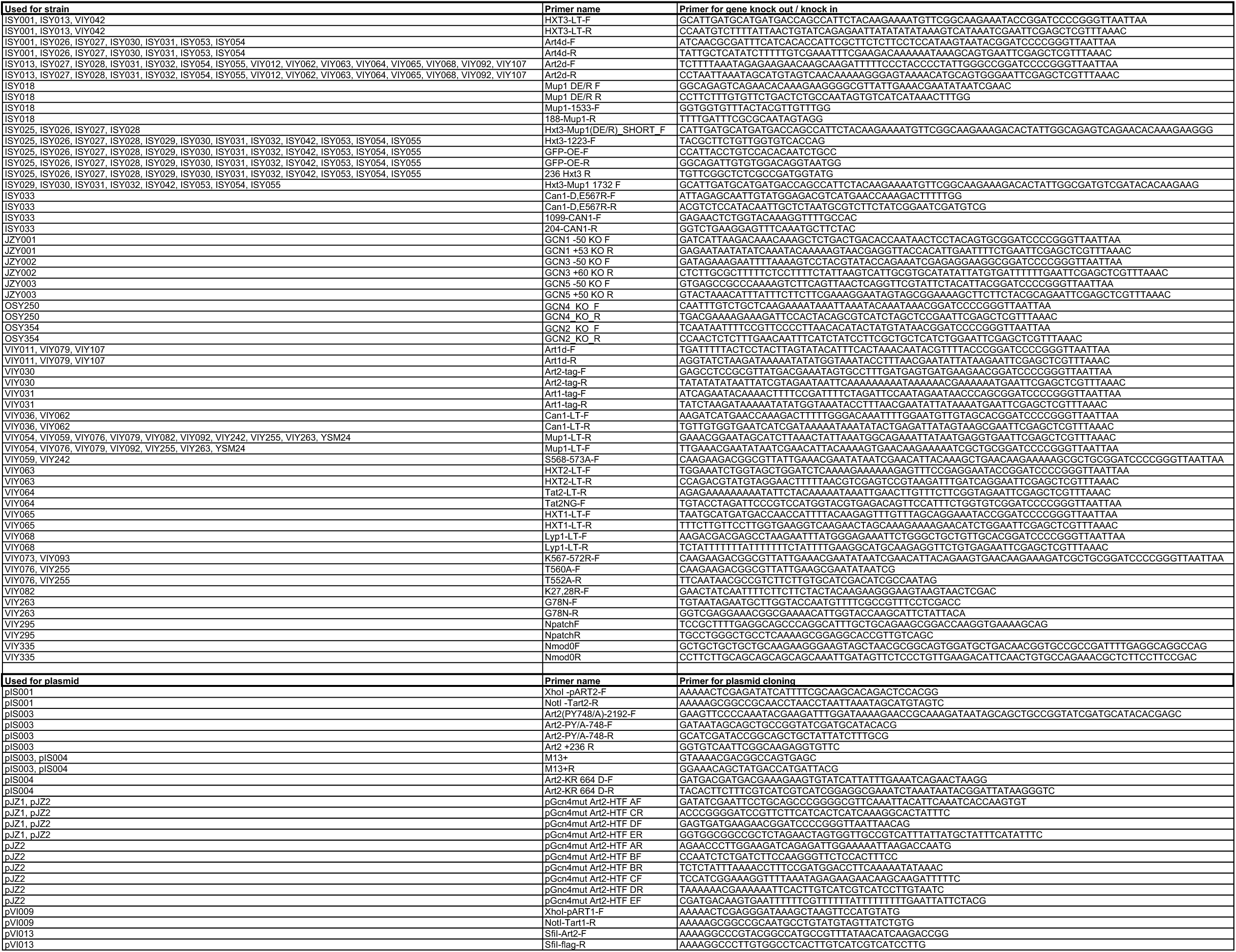
Yeast strains, plasmids and primer.

